# Discovery of the rosalexin pathway expands the modular network of maize diterpenoid chemical defenses

**DOI:** 10.64898/2026.04.15.718773

**Authors:** Anna E. Cowie, Gabrielle Wyatt, Siena Schumaker, Ahmed Khalil, Alicia Ross, Shamita Bhattacharjee, Jishnu Narayanan, Yezhang Ding, David Hurd, Jedediah Peek, Elly Poretsky, Alisa Huffaker, Dean J. Tantillo, Dan T. Major, Eric A. Schmelz, Philipp Zerbe

**Affiliations:** Department of Plant Biology, University of California-Davis, Davis, CA 95616, USA; Section of Cell and Developmental Biology, University of California-San Diego, La Jolla, CA 92093, USA; Department of Chemistry, University of California-Davis, Davis, CA 95616, USA; Department of Chemistry and Institute for Nanotechnology and Advanced Materials, Bar Ilan University, Ramat Gan 5290002, Israel

**Keywords:** Zea mays, terpenoids, diterpene synthases, cytochrome P450 monooxygenases, phytoalexins, plant immunity, plant specialized metabolism

## Abstract

The evolutionary expansion of specialized metabolism has shaped the ability of plants to adapt to combined pathogen, pest, and other environmental pressures. For instance, the duplication and divergence of ancestral gibberellin pathway genes have given rise to specialized kauralexin and dolabralexin diterpenoids in maize (*Zea mays*) that serve as core components of disease resistance and stress adaptation. Here, we describe the biosynthesis and elicited production of rosalexins as a previously unrecognized component of the maize chemical defense network. By integrating genomics-enabled gene discovery, combinatorial enzyme assays, and AI-assisted enzyme mechanistic studies we show that maize rosalexin biosynthesis proceeds via a distinct 5-rosanol scaffold formed by the pairwise activity of two diterpene synthases, ZmTPS38/CPS2/AN2 and ZmTPS42/KSL1, recruited from gibberellin metabolism. Further oxygenation by the promiscuous P450 enzyme, ZmCYP71Z18, yields epoxyrosanol that, in turn, can undergo epoxide ring opening to form trihydroxyrosanol. Epoxyrosanol, but not 5-rosanol or trihydroxyrosanol, display strong inhibitory activity on fungal pathogen growth *in vitro*, highlighting the contribution of the epoxide group to antibiotic efficacy. Large variation in rosalexin presence and abundance exists across maize genotypes due to expansive *ZmTPS42/KSL1* gene sequence variation and pseudogenization. Transcriptomics and targeted metabolomics demonstrated the pathogen-elicited accumulation of rosalexins in maize lines featuring functional *ZmTPS42/KSL1* genes. However, no dominant pathogen resistance phenotype was observed in association with rosalexin abundance. These collective findings expand our knowledge of how multiple interconnected diterpenoid pathways arose in maize via duplication of hormone-metabolic genes and enable the utilization of a common precursor to form modular chemical defense layers.

**Significance Statement:** Plant diterpenoids play critical roles in crop development, stress defense and ecological adaptation. In maize, diterpenoids serve as key components of chemical defenses against pests and diseases with direct impact on crop immunity and vigor. Enzymes of the diterpene synthase and cytochrome P450 families largely drive diterpenoid chemical diversity. This study reports the discovery and characterization of the pathway forming rosalexin diterpenoids in maize. Pathogen-elicited accumulation and in vitro antifungal activity of rosalexin metabolites supports a physiological function in maize chemical defense.

## Introduction

Plants employ complex blends of specialized metabolites to mediate interactions with their environment and thrive in various ecological niches ^1,2^. Among these metabolites, the large group of 20-carbon diterpenoids serves multi-facetted functions with direct impact on plant vigor and fitness ^3^. Widely conserved diterpenoids such as gibberellin (GA) phytohormones play critical roles in plant development. By contrast, specialized diterpenoids are narrowly distributed across plant lineages, often display stimulus- and tissue-specific accumulation, and function in mediating the outcome of defensive and cooperative plant-environment interactions ^3,4^. For example, specialized diterpenoids serve as core chemical defenses against microbial diseases, insect pests and abiotic stressors in major grain crops such as maize (*Zea mays*) and rice (*Oryza sativa*) ^5,6^. Diterpenoid pathways with demonstrated or predicted functions in stress defense also exist in wheat (*Triticum aestivum*) ^7^, millet (*Setaria italica*) ^8^, barley (*Hordeum vulgare*) ^9^, and switchgrass (*Panicum virgatum*) ^10,11^, illustrating the broad distribution of species-specific networks that produce bioactive diterpenoids in monocot crops. Driven by environmental selection pressures, this expansion of diterpenoid metabolism has been enabled by the repeated duplication and functional divergence of ancestral pathway genes of general metabolism ^1,12,13^. In particular, GA-biosynthetic diterpene synthases (diTPSs) and cytochrome P450 monooxygenases (P450s) have provided a genetic reservoir for the duplication and functional evolution of specialized enzyme functions ^12,14^. The resulting diTPS and P450 gene families comprised of functionally distinct members form modular pathway networks, where different combinations of class II and class I diTPSs and P450s generate various structural scaffolds and functional decorations that yield diterpenoids with specific physiological functions ^3,14,15^. In maize, this modular metabolic network generates unique groups of protective diterpenoids, including kauralexins and dolabralexins ^6,16–18^ (**Fig. 1**). Kauralexins exhibit strong antimicrobial and insecticidal activities, and accumulate locally at the site of attack predominantly in above-ground tissues ^16,19,20^. Dolabralexins are largely root-specific, formed in response to both biotic and abiotic stress, and may contribute to rhizosphere microbiome interactions ^16,17,21–23^. Due to their evolutionary relationship, GAs, kauralexins and dolabralexins share a common *ent*-copalyl diphosphate (*ent*-CPP) precursor ^12,16^ (**Fig. 1**). Dysregulation of GA and defensive diterpenoid pathways in maize is mitigated by the presence of two duplicated class II diTPSs, ZmTPS37/CPS1/ZmAN1 and ZmTPS38/CPS2/ZmAN2, that convert the common geranylgeranyl diphosphate (GGPP) substrate into *ent*-CPP, yet display distinct spatiotemporal regulation and stress-inducibility. While ZmTPS37/CPS1/ZmAN1 functions in GA metabolism ^24^, ZmTPS38/CPS2/ZmAN2 is stress-inducible and involved in specialized diterpenoid metabolism ^19,25^. Downstream, two class I diTPSs, namely ZmTPS43/KSL2 and ZmTPS45/KSL4, control key pathway nodes by converting *ent*-CPP into the kauralexin precursor *ent*-isokaurene or the dolabralexin precursor dolabradiene, respectively ^16,17,21^. Catalytically promiscuous P450s of the CYP701A and CYP71Z subfamilies that are capable of functionally modifying both precursors then bring about the biosynthesis of different, bioactive kauralexins and dolabralexins ^16,17^. Presence of two additional class II diTPSs, ZmTPS39/CPS3 and ZmTPS40/CPS4 that form (+)-CPP and 8,13-CPP, respectively, further shows that the diversity of maize specialized diterpenoids extends beyond kauralexins and dolabralexins ^18^.

**Fig. 1.**
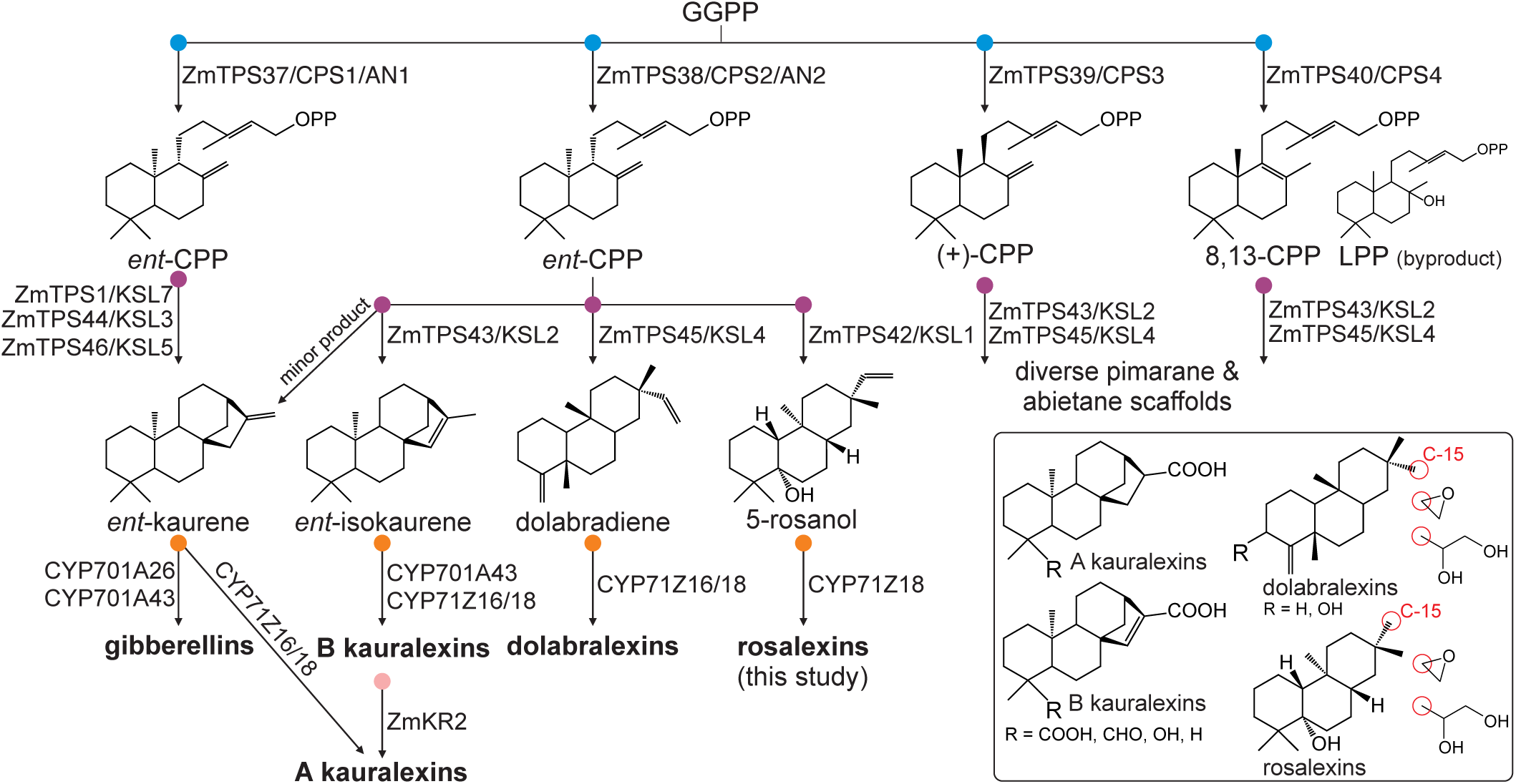
Schematic overview of the maize diterpenoid-biosynthetic network. Maize contains a modular diterpenoid-metabolic network to form specialized compound groups with distinct bioactivities, including A- and B-type kauralexins, dolabralexins, and rosalexins (this study). Abbreviations: GGPP, geranylgeranyl diphosphate; CPP; copalyl diphosphate; CPS, copalyl diphosphate synthase; KSL, kaurene synthase-like.

Here, we report that the maize class I diTPS, ZmTPS42/KSL1, catalyzes a previously unrecognized reaction that transforms *ent*-CPP into a 5-rosanol scaffold. Modular activity of ZmTPS38/CPS2/ZmAN2, ZmTPS42/KSL1, and the promiscuous P450 enzyme, ZmCYP71Z18, form a third *ent*-CPP-derived pathway branch in maize en route to specialized diterpenoids, coined here rosalexins. *In vitro* antimicrobial activity of epoxyrosanol along with pathogen-induced *ZmTPS42/KSL1* transcript and rosalexin accumulation in aerial tissues indicated a defense-related function. However, no clear pathogen-susceptible phenotype was observed in *ZmTPS42/KSL1*-deficient maize plants, suggesting a more specialized stress-responsive function within the network of bioactive kauralexin, dolabralexin, and rosalexin diterpenoids in maize.

## Results

### ZmTPS42/KSL1 shows high sequence variation in maize

Unlike related grain crops such as rice, wheat, and barley, where gene clusters for species-specific specialized diterpenoid pathways have been identified ^7,26^, gene loci of maize diterpenoid pathways, including known GA, kauralexin, and dolabralexin biosynthesis are broadly dispersed across the maize genome. Mining of the maize B73 genome (Zm-B73-REFERENCE-NAM-5.0) located *ZmTPS42/KSL1* (*Zm00001eb176190*) on chromosome 4 (chr4:55755497-55756759) within a distance of 48.7 Mb to the next known terpenoid-metabolic gene, i.e. the labda-8,13-dien-15-yl diphosphate synthase, *ZmTPS40/CPS4* ^18^ (**Fig. 2a**). However, closer inspection revealed that *ZmTPS42/KSL1* in the B73 line lacks the *γ*- and β-domains as well as the catalytic DDxxD motif and, thus, likely represents a pseudogene (**Fig. 2b**). We therefore examined the *ZmTPS42/KSL1* sequences across 13 selected inbred lines, including founder lines of the Nested Association Mapping (NAM) population and the Goodman diversity panel, which revealed substantial sequence variations in the *ZmTPS42/KSL1* gene (**Fig. 2b, Supplementary Fig. S1**). Similar to B73, several lines including Mo17, CML133, B73, B75, and W22, featured major sequence variations and gaps, predictably rendering the encoded proteins non-functional. By contrast, Oh7B, EP1, CML52, CML69, and CML277 among other lines contain full-length *ZmTPS42/KSL1* genes with predicted protein sequence identities of 74-100%. Using ZmTPS42/KSL1 from the maize line Oh7B as a representative full-length protein, phylogenetic analysis placed ZmTPS42/KSL1 separate from known *ent*-kaurene synthases involved in GA metabolism and within a group of class I diTPSs that include the *ent*-iso-kaurene synthase, ZmTPS43/KSL2, the dolabradiene synthase, ZmTPS45/KSL4, as well as specialized class I diTPSs from switchgrass, thus suggesting a related function in specialized metabolism (**Fig. 2c**).

**Fig. 2.**
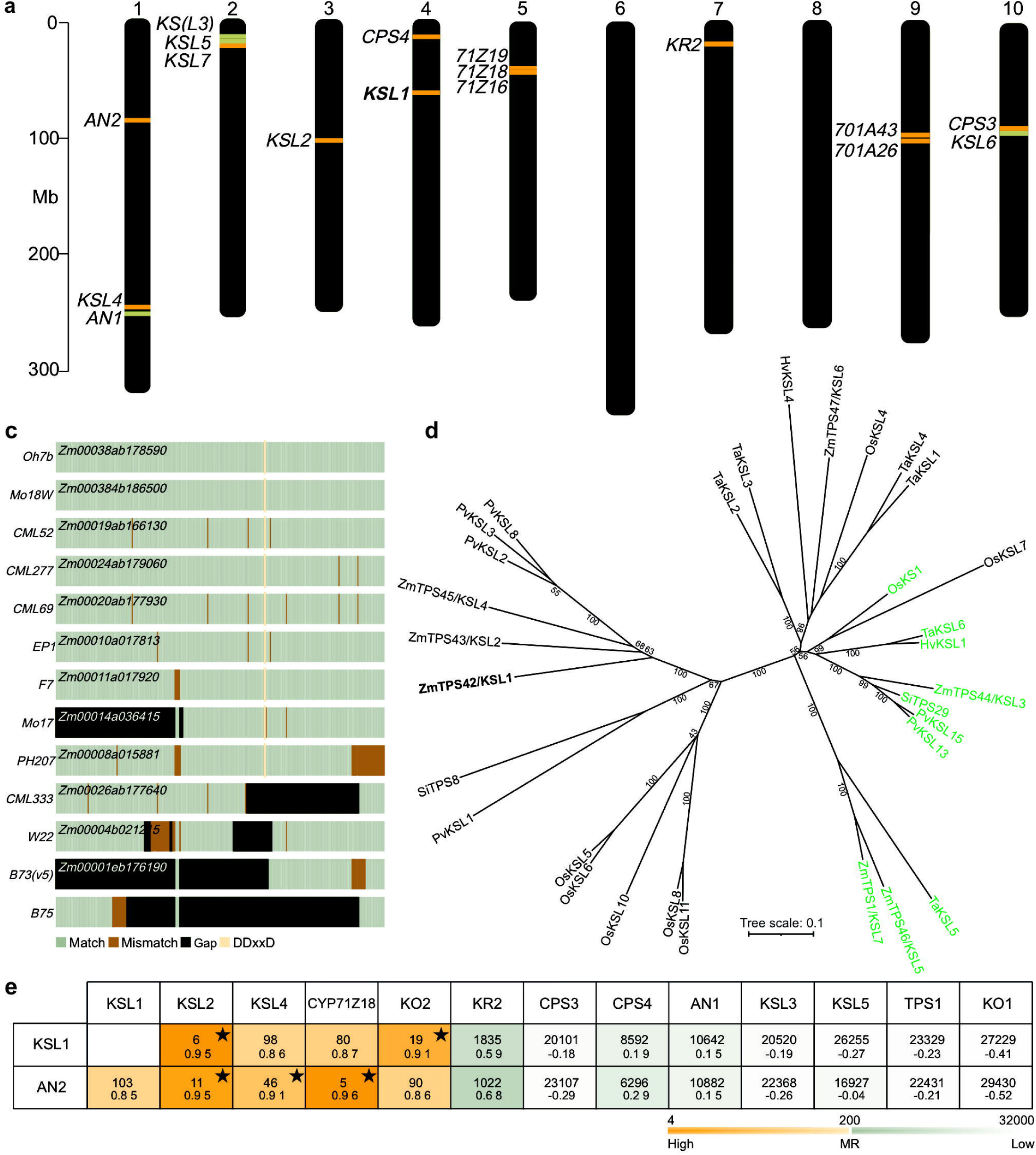
*Zm*TPS42/KSL1 is a diterpene synthase with high sequence variation across maize genotypes. **a**, Chromosome plot (Zm-B73-REFERENCE-NAM-5.0) depicting genome locations of diterpenoid-metabolic genes. Genes associated with gibberellin metabolism (green) or specialized diterpenoid metabolism (orange) are highlighted. **b**, Protein sequence alignment of ZmTPS42/KSL1 sequences derived from selected maize lines. **c**, Unrooted maximum likelihood phylogenetic tree (500 bootstrap repetitions) of ZmTPS42/KSL1 from the Oh7B line and biochemically characterized monocot class I diTPSs. Known GA-metabolic enzymes are depicted in green. **d**, Heat map illustrating the pairwise gene correlation for *ZmTPS38/CPS2/AN2* and *ZmTPS42/KSL1* genes with class II diTPS and GA metabolism genes. Strong co-expression was defined by Pearson’s correlation >0.7 and mutual rank (MR) of <200). Complementary Pearson’s correlation coefficient (PCC) of >0.7 and mutual rank value (MR) of <200 are shaded in orange (using -log10(MR)), and weak correlations in grey. PCC and MR values are printed on bottom and top respectively. Stars denote gene pairs with PCC>0.8 and MR <50. Abbreviations: CPS, copalyl diphosphate synthase; KSL, kaurene synthase-like; CYP, cytochrome P450; KO, kaurene oxidase; KR, kauralexin reductase; Ta, *Triticum aestivum*; Hv, *Hordeum vulgare*; Zm, *Zea mays*; Os, *Oryza sativa*; Pv, *Panicum virgatum*; Si, *Setaria italica*.

Gene expression profiles available in public repositories including the Sekhon gene expression atlas showed highest expression of *ZmTPS42/KSL1* in above-ground tissues including stem internodes, first and post-pollination leaves ^27,28^ as well as embryo tissue ^29^. In addition, integrating mutual rank (MR) and Pearson’s correlation (PCC) analyses of in-house transcriptome data of maize stems infected with the major maize pathogen *Fusarium verticillioides* showed patterns of co-expression of *ZmTPS42/ZmKSL1* and known genes of maize specialized diterpenoid metabolism (**Fig. 1d, Supplementary Table S1**). Specifically, *ZmTPS42/ZmKSL1* gene expression was strongly correlated with *ZmTPS43/KSL2, ZmTPS45/KSL4, ZmCYP71Z18,* and *CYP701A43/KO2* involved in kauralexin and dolabralexin metabolism. Notably, among the four maize class II diTPSs, *ZmTPS42/ZmKSL1* showed the strongest co-regulation with the *ent*-CPP synthase *ZmTPS38/CPS2/ZmAN2*. By contrast, GA-metabolic genes, including *ZmTPS37/CPS1/AN1*, *ZmTPS44/KS(L3)/D5*, and *ZmTPS46/KSL5* showed distinct expression patterns from *ZmTPS42/ZmKSL1* (**Fig. 2d**).

### ZmTPS42/KSL1 forms a rosane-type diterpene scaffold

To test the biochemical function of ZmTPS42/KSL1 in the context of the maize diterpenoid network, we selected the full-length ZmTPS42/KSL1 from Oh7B to assay pairwise enzyme activities with known maize class II diTPSs: the *ent-*CPP synthase ZmTPS38/CPS2/AN2 (catalytically identical to ZmTPS37/CPS1/AN1) ^25^, the (+)-CPP synthase ZmTPS39/CPS3, and the 8,13-CPP synthase ZmTPS40/CPS4 ^18^. *Agrobacterium*-mediated transient co-expression assays in *Nicotiana benthamiana* were conducted by pairing ZmTPS42/KSL1 individually with each class II diTPS. Co-expression of ZmTPS42/KSL1 with *Zm*TPS38/CPS2/AN2 resulted in the formation of a major product (compound **1**) and a minor byproduct (compound **2**). Compound **1** featured *m*/*z* 275 and *m*/*z* 290 mass ions indicative of a labdane-type diterpene alcohol and product **2** featured *m/z* 257 and *m/z* 272 mass ions characteristic of a labdane-type olefin (**Fig. 3a-b**). In addition, GC-MS analysis of the extracted enzyme products of the pairwise activity of ZmTPS42/KSL1 with the (+)-CPP synthase ZmTPS39/CPS3 showed several labdane diterpenoids (compounds **3-8**), albeit at very low abundance. Among these, **3**, **6**, **7,** and **8** were distinct ZmTPS42/KSL1 products and **3** could be identified as sandaracopimaradiene based on comparison to an authentic standard (**Fig. 3a, Supplementary Fig. S2**). Combined activity of ZmTPS42/KSL1 with the 8,13-CPP synthase ZmTPS40/CPS4 yielded a range of unidentified labdane scaffolds (compounds **9-12**) at low abundance (**Fig. 3a, Supplementary Fig. S2**). However, these products represented byproducts of ZmTPS40/CPS4 rather than unique compounds resulting from ZmTPS42/KSL1 activity.

**Fig. 3.**
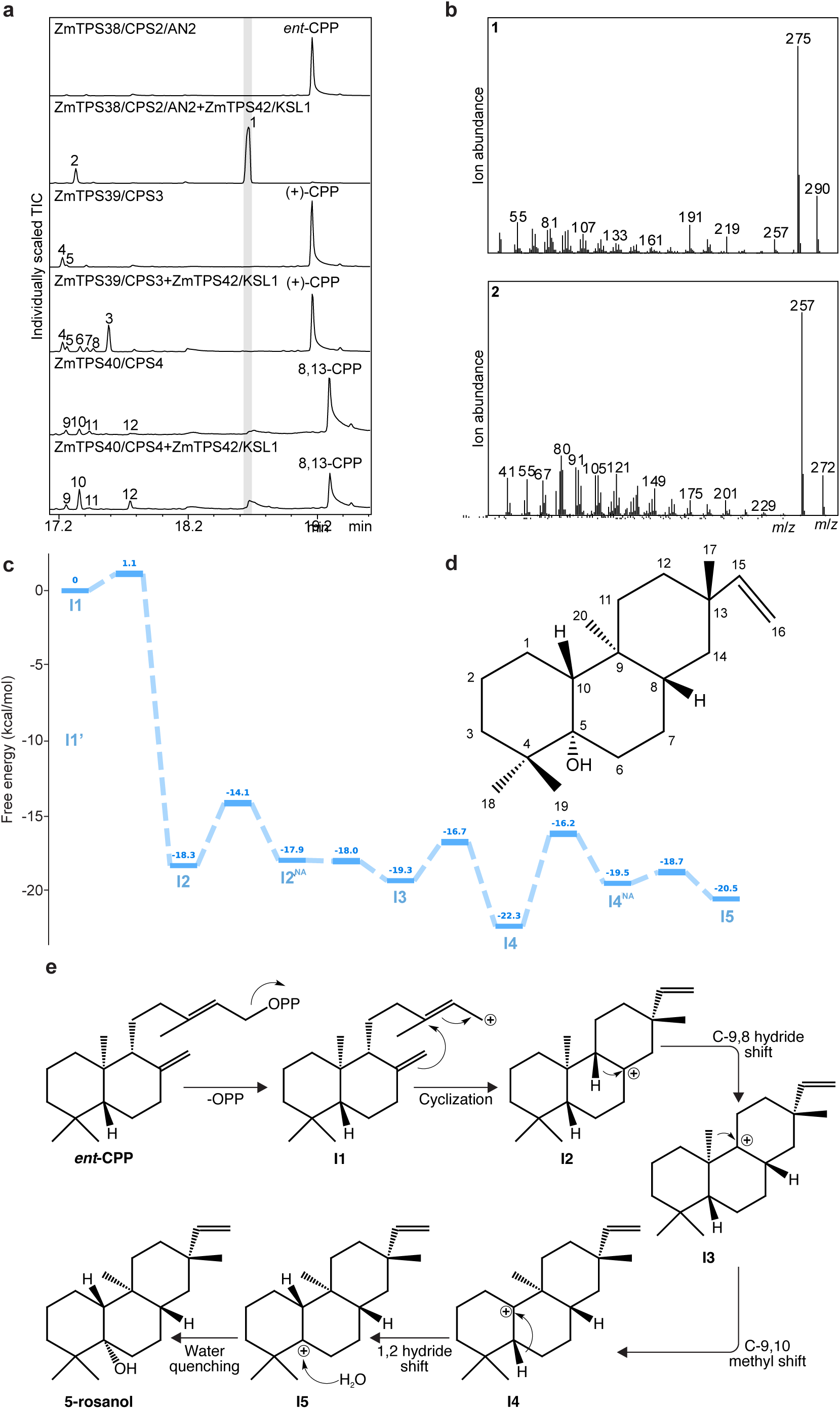
Functional characterization of *Zm*TPS42/KSL1. **a**, Total ion chromatograms (TIC) of reaction products resulting from *Agrobacterium*-mediated *Nicotiana benthamiana* co-expression assays of ZmTPS42/KSL1 with ZmTPS38CPS2/AN2, ZmTPS39/CPS3, or ZmTPS40/CPS4. **b**, Mass spectrum of compound 1 and 2 resulting from the pairwise reaction of ZmTPS38/CPS2/AN2 and ZmTPS42/KSL1. **c**, Free energy diagram of the proposed ZmTPS42/KSL1 reaction mechanism leading to the final intermediate at M06-2X/def2-SVP level of theory. **d**, Structural identification of compound **1** as 5-rosanol, as verified by NMR analysis. **e**, Proposed ZmTPS42/KSL1-catalyzed reaction mechanism of the conversion of *ent*-CPP into 5-rosanol.

Given that the highest ZmTPS42/KSL1 product abundance was observed with *ent*-CPP as a substrate and the strong co-regulation patterns of *ZmTPS38/CPS2/AN2* and *ZmTPS42/KSL1* (**Fig. 2d**), we investigated this reaction further. To elucidate the structure of the ZmTPS42/KSL1 product **1** we enzymatically produced and purified >1 mg of **1** for NMR analysis and identified the metabolite as 5-hydroxy-rosa-15-ene (hereafter named 5-rosanol) (**Fig. 3d, Supplementary Fig. S3**). Complementary to experimental NMR data, we employed computational NMR analysis of 16 possible isomers across the four stereocenters present in 5-rosanol to assign absolute stereochemistry. Experimental 1D (^1^H and ^13^C) and 2D (H2BC, HMBC, COSY, NOESY) NMR data assessed with complementary DP4+ calculations matched compound **1** to stereoisomer 11, identifying the ZmTPS42/KSL1 product 5*S*,8*S*,9*S*,10*R*,13*S*)-5-hydroxy-rosa-15-ene (**Fig. 3c; Supplementary Tables S2 & S3**) ^30^. Verification of the 5-rosanol structure next enabled us to apply the AI-assisted automated reaction mechanism discovery platform, RxnNet ^31^ to propose a catalytic mechanism of 5-rosanol biosynthesis by ZmTPS42/KSL1 (**Fig. 3c,e**). Following abstraction of the pyrophosphate group of *ent*-CPP, the reaction proceeds via cyclization of **I1** (set as the reference state with a free energy of 0.0 kcal/mol for its global minimum conformer in the gas-phase) to form the pimara-15-en-8-yl^+^ carbocation (**I2**). Notably, a slight 1.1 kcal/mol free energy barrier arises from a conformational change, which is followed by a barrierless cyclization process. A subsequent ring-puckering of the global minimum conformer of **I2** (ΔG^‡^ = 4.1 kcal/mol) results in the near-attack conformer (NAC) ^32,33^ of **I2** (**I2^NA^**). The 1,2-hydride shift between **I2^NA^** and **I3** is a barrierless process, as the latter is already in a NAC state ready for the 1,2-hydride shift. The **I3** to **I4** methyl migration proceeds via a free energy barrier of 2.6 kcal/mol. The subsequent hydride shift goes via an initial conformational change to reach **I4^NAC^** with a ΔG^‡^ = 6.1 kcal/mol followed by the chemical step which has a barrier of 0.8 kcal/mol. Finally, water quenching of the carbocation center of **I5** results in 5-rosanol (**Supplementary Fig. S4 & Table S4**). Intermediates **I2** through **I5** exhibit similar free energies and are characterized by low-barrier carbocationic transformations. Consequently, an equilibrium may exist among these species, with the final product distribution being primarily determined by the positioning of the active-site water, which is analogous to that reported for active-site base in taxadiene synthase ^34–36^. Low product abundance prevented NMR analysis of the ZmTPS42/KSL1 byproduct **2**. However, based on characteristic *m*/*z* 257 and *m*/*z* 272 mass ions and the biosynthetic mechanism forming 5-rosanol, we hypothesize that **2** represents a rosadiene olefin scaffold derived from neutralization of the intermediary rosa-15–en-10-yl^+^ (**I4**) or rosa-15-en-5-yl^+^ (**I5**) carbocations via deprotonation rather than water quenching ^37^.

Given that the biosynthesis of related kauralexin and dolabralexin diterpenoids requires the functional modification of the respective diTPS-produced scaffolds by specialized P450 enzymes of the CYP701A and CYP71Z subfamilies ^16,17,38^, we performed co-expression studies combining ZmTPS38/CPS2/AN2 and ZmTPS42/KSL1 (Oh7B) with known diterpenoid-metabolic maize P450s. Co-expression with ZmCYP71Z18 yielded a new major (compound **13**) and minor (compound **14**) product, featuring mass ions of *m*/*z* 306 and *m*/*z* 288, respectively, that are indicative of oxygenated labdane diterpenoids (**Fig. 4a**). Purification and NMR analysis of **13** and **14** verified these structures as 5-hydroxy-15-epoxyrosa-5(6)-ene (hereafter named epoxyrosanol) and epoxy-rosa-5(6)-ene (named epoxyrosaene), respectively (**Fig. 4b, Supplementary Figs. S5 & S6**). While epoxyrosanol is derived from 5-rosanol, the formation of small quantities of epoxyrosaene supported our hypothesis that the minor ZmTPS42/KSL1 product **2** is rosaene and serves as a precursor for epoxyrosaene. In keeping with the nomenclature of other maize diterpenoids, we collectively designated 5-rosanol (**1**), rosaene (**2**), epoxyrosanol (**13**), and epoxyrosaene (**14**) as rosalexins. Functional testing of ZmTPS38/CPS2/AN2 and ZmTPS42/KSL1 with other characterized maize diterpenoid-metabolic P450s, including ZmCYP71Z16 ^17^, ZmCYP71Z19 ^38^, ZmCYP701A26 ^16,39^, and ZmCYP701A43 ^16^, yielded no additional products (**Supplementary Fig. S7**).

**Fig. 4.**
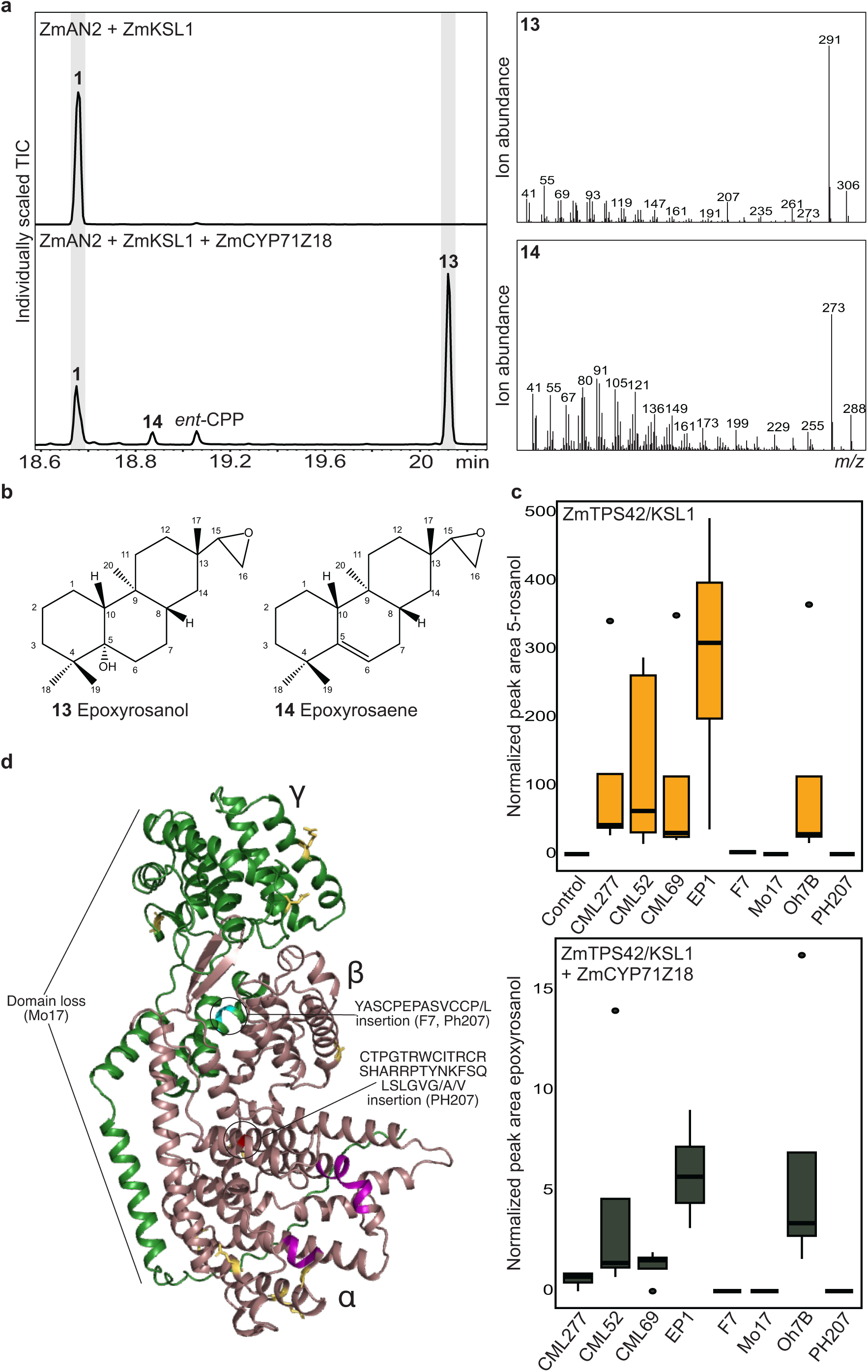
Identification of the rosalexin pathway. **a**, Total ion chromatograms (left) and mass spectra (right) of reaction products resulting from *Nicotiana benthamiana* co-expression assays of ZmTPS38/CPS2/AN2 and ZmTPS42/KSL1 only, or with the addition of ZmCYP71Z18. **b**, Structural identification via NMR analysis of compounds **13** and **14** as epoxyrosanol and epoxyrosaene, respectively. **c**, Normalized abundance of 5-rosanol (top) and epoxyrosanol (bottom) produced by ZmTPS42/KSL1 from each maize line co-expressed with *Zm*TPS38/CPS2/AN2 for 5-rosanol and ZmTPS38/CPS2/AN2 and ZmCYP71Z18 for epoxyrosanol. Error bars represent standard error of the mean (n=4). **d**, AlphaFold 3 homology model of Oh7B ZmTPS42/KSL1 (brown) with the DDxxD and NSE/DTE motifs highlighted in magenta. Sequence variations in ZmTPS42/KSL1 sequences derived from other tested lines are highlighted in yellow.

Having established the enzymatic function of ZmTPS42/KSL1 derived from the Oh7B maize line, we further assessed the activity of predicted ZmTPS42/KSL1 proteins from other maize lines using co-expression assays with ZmTPS38/CPS2/AN2 alone or with ZmTPS38/CPS2/AN2 and ZmCYP71Z18. The partial protein sequence of Mo17 ZmTPS42/KSL1, lacking a substantial segment of the N-terminal *γβ*-domain, expectedly resulted in an inactive enzyme (**Fig. 4c-d**). Likewise, the full-length ZmTPS42/KSL1 proteins derived from the maize lines F7 and PH207 did show no or only trace enzyme activity, likely as a result of more extensive sequence variation and insertions (**Fig. 4c-d**). By contrast, rosalexin production was observed for the full-length ZmTPS42/KSL1 proteins derived from the CML277, CML52, CML69, Oh7B, and EP1 maize lines (**Fig. 4c-d**). Notably, 5-rosanol product levels varied significantly (p = 0.036) across ZmTPS42/KSL1 proteins from these tested maize genotypes.

### Stress-elicited rosalexin accumulation in planta reveals additional pathway products

To investigate the presence of rosalexins *in planta*, we performed targeted metabolomics of maize stem and root tissues of 53-day-old plants using contrasting maize lines containing inactive (Mo17) or functional (Oh7B) *ZmTPS42/KSL1* genes three days after pathogen infection with *F. verticillioides* as compared to mock-treated (wounding only) control samples. Consistent with the absence of a functional *ZmTPS42/KSL1*, LC-MS/MS analysis via quantifying relative ion intensity and abundance showed that Mo17 tissues did not contain rosalexin metabolites at any detectable level irrespective of pathogen or mock treatment (**Fig. 5; Supplementary Tables 5 & 6**). Control Oh7B plants also did not contain detectable rosalexin levels. However, accumulation of 5-rosanol and epoxyrosanol was observed in pathogen-infected roots and stems of Oh7B plants. Abundance of all compounds was 10-fold higher in stems as compared to roots consistent with lower *ZmTPS42/KSL1* gene expression in roots (**Fig. 5a-b**). Standard-verified 5-rosanol was detected by a predominant *m*/*z* 273 mass fragment representing a neutral loss of water [M-H_2_O+H]^+^. On average (n=4), infected plants accumulated 5-rosanol at 4.3 ± 1.1 μg g^-1^ (FW) in roots and at 59.5 ± 14.6 μg g^-1^ (FW) in stems. Ion spectra analysis of epoxyrosanol revealed a *m*/*z* 307 fragment [M+H]^+^ consistent with the NMR molecular formula of the purified compound, as well as a predominant *m*/*z* 330 ion representing a sodium adduct [M+Na]^+^. Based on these ions, epoxyrosanol was detected exclusively in infected Oh7B stems and roots and at trace amounts concentrations of 0.76 ± 0.38 ng g^-1^ (FW) and 0.015 ± 0.015 ng g^-1^ (FW), respectively (**Fig. 5a-b**). In infected Oh7B stem tissue, another abundant metabolite (compound **15**) was detected, which also featured a *m/z* of 307 mass fragment and a near identical MS^2^ fragmentation pattern as compared to expoxyrosanol, yet showed an earlier retention time of 2.7 min as compared to epoxyrosanol at 5.5 min (**Fig. 5c**). Our prior work had demonstrated the presence of trihydroxydolabrene (THD) as a major metabolite derived from the dolabralexin pathway in roots of field-grown maize ^17^. We therefore hypothesized that compound **15** could be a similar structure, resulting from the enzymatic or spontaneous opening of the epoxide ring of epoxyrosanol. We tested this hypothesis using a chemical conversion approach. Subjecting enzymatically produced, purified epoxyrosanol to slightly acidic conditions indeed yielded **15** (**Fig. 4e**). Further NMR analysis indeed identified **15** as trihydroxyrosanol (THR), adding another compound to the group of rosalexins that features the loss of the epoxide and gain of two hydroxyl groups at C-15 and C- 16 (**Fig. 5d, Supplementary Fig. S8**). Re-analyzing pathogen-infected plant material demonstrated the presence of THR in infected roots, albeit at trace amounts. Conversely, THR significantly accumulated in stem tissue at 5.3 μg g^-1^ (FW) (**Fig 5a-b**). No THR was detected in control plants.

**Fig. 5.**
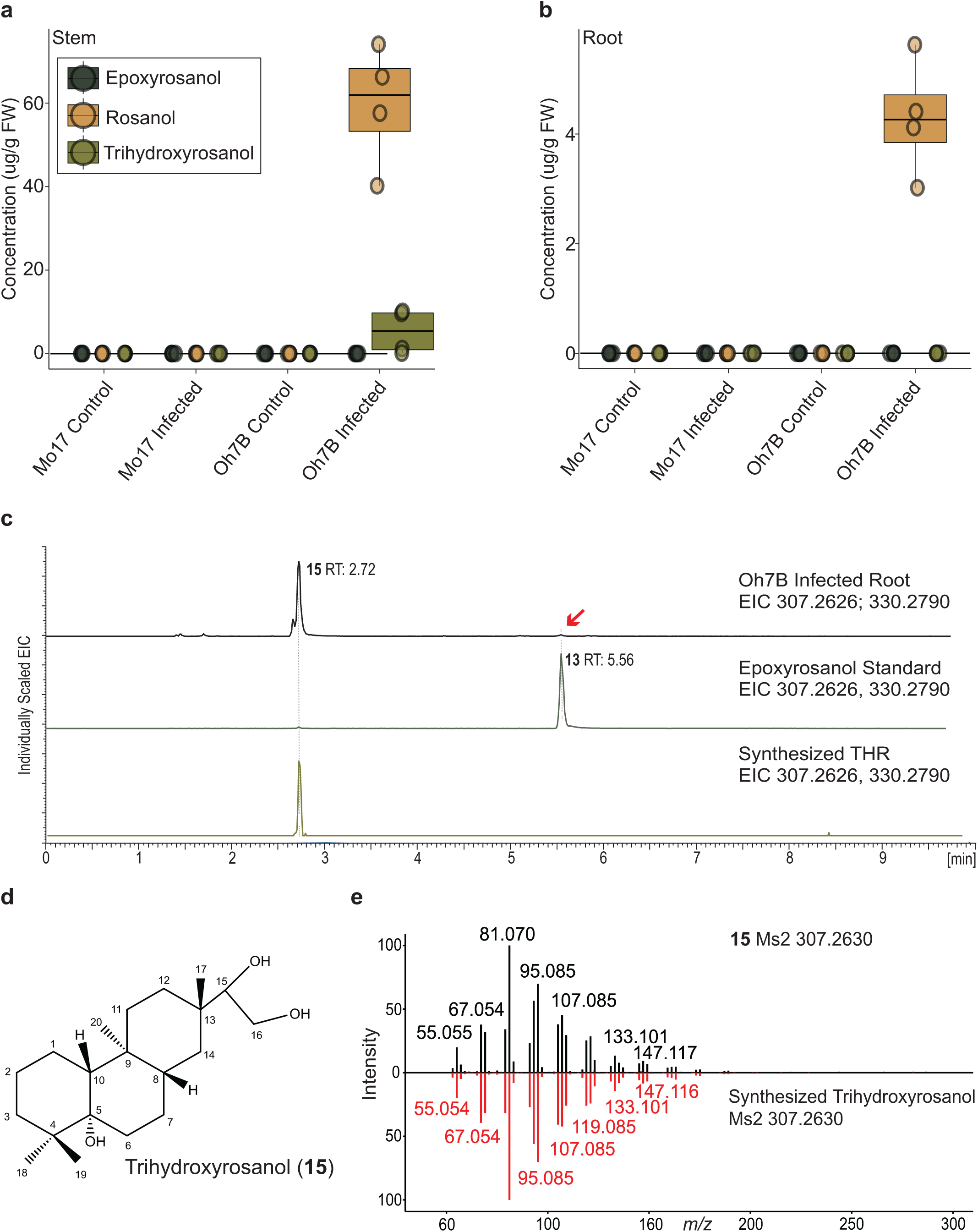
Rosalexins accumulate *in planta* in response to pathogen infection. **a-b**, LC-MS/MS average metabolite abundance in stem (**a**) and root (**b**) for epoxyrosanol, 5-rosanol and trihydroxyrosanol (THR) in Mo17 and Oh7B plants in mock-treated (control) and plants infected with *Fusarium verticillioides* (infected). Boxes represent the interquartile range (IQR), center line indicates the median, and whiskers extend to 1.5×IQR (n=4). Raw metabolite abundance data are given in **Supplementary Tables S5 & S6**. **c,** LC-MS/MS extracted ion chromatogram (EIC) at 307.2626 and 330.2790 *m/z* of infected Oh7B roots, purified epoxyrosanol standard and synthesized trihydroxyrosanol (THR) standard. **d**, NMR verified structure of synthesized THR. **e,** LC-MS2 spectra comparing *in planta* compound **15** (black) and synthesized THR standard (red).

### Rosalexins display antibiotic activity against fungal pathogens

To assess the antifungal potential of rosalexins, we performed *in vitro* fungal growth inhibition assays using the maize pathogens *F. verticillioides*, *Aspergillus parasiticus*, and *Rhizopus microsporus*, which cause maize seedling blight, stalk, ear and root rots, and mycotoxin contamination. Fungal growth was measured by optical density at 600 nm (OD_600_) in the presence of purified 5-rosanol, epoxyrosanol, and THR. Following prior bioactivity assays of maize kauralexins and dolabralexins, we utilized concentrations of 10, 25, and 50 μg mL^-1^ for each metabolite to test for antimicrobial activity ^16,17^. With respect to *F. verticillioides*, 5-rosanol displayed no significant growth-inhibitory activity irrespective of compound concentration (**Fig. 6, Supplementary Table S7**). In the presence of THR, a brief increase in fungal growth was observed, while a moderate inhibitory effect was detected at the 48 h end point. By contrast, epoxyrosanol significantly inhibited fungal growth at all tested compound concentrations. A similar impact on fungal growth was observed for *A. parasiticus* with epoxyrosanol showing the strongest growth-inhibitory activity. By contrast, 5-rosanol and THR had a strong inhibitory effect on the growth of *R. microsporus* for ∼32 h, after which fungal growth rapidly increased to near-control levels. Similar to *F. verticillioides* and *A. parasiticus*, epoxyrosanol showed strong inhibitory activity against *R. microsporus*. For example, at a concentration of 25 μg mL^-1^, epoxyrosanol inhibited *R. microsporus* by 2.43 fold (p < 0.000000001), as compared to 1.96 fold (p = 0.000000455) for *F. verticillioides*, and 1.53 fold (p = 0.000149473) for *A. parasiticus* (**Fig. 6, Supplementary Table S7**).

**Fig. 6.**
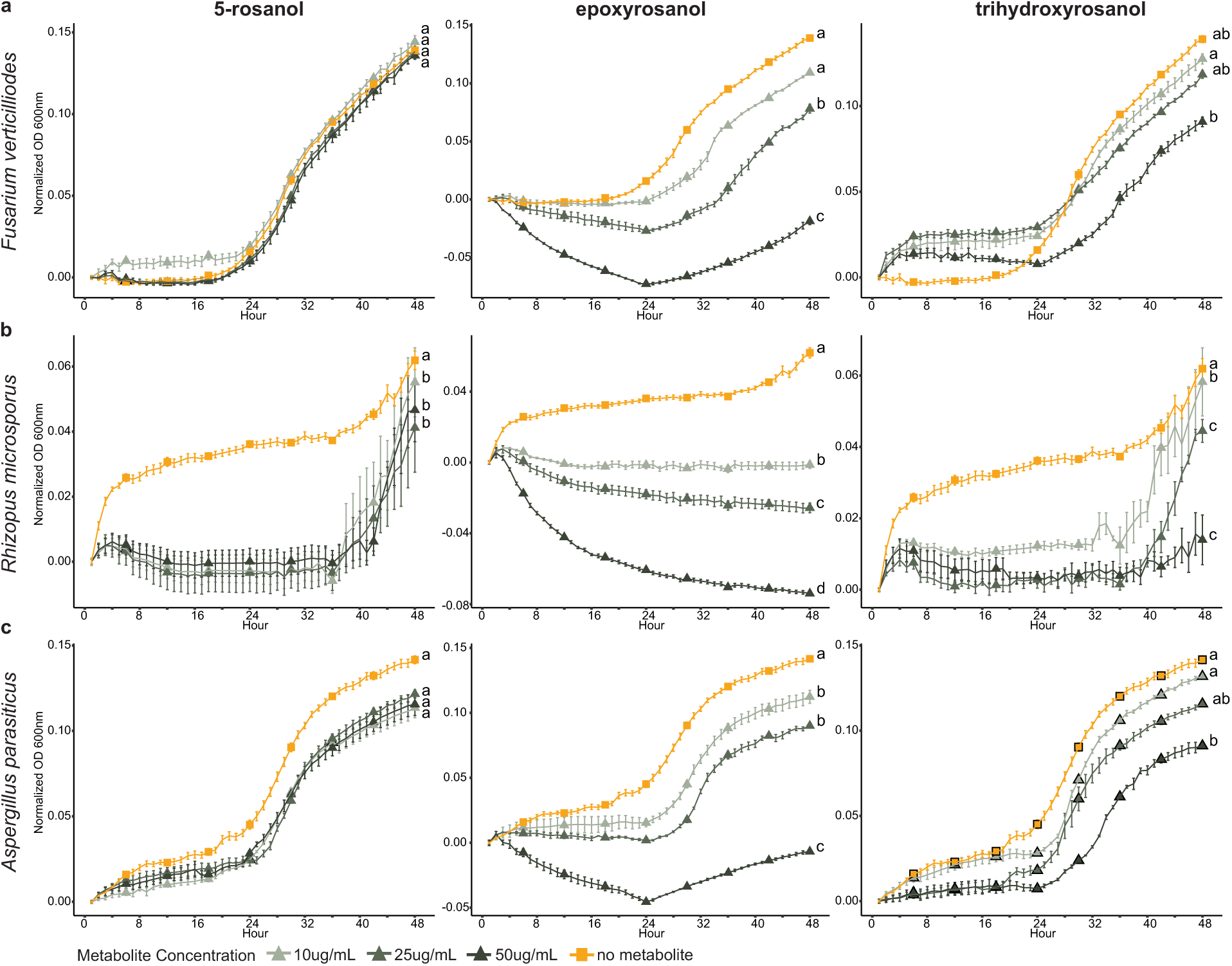
Antifungal bioactivity of rosalexins. Average growth (OD_600_) of *Fusarium verticillioides* (**a**)*, Rhizopus microsporus* (**b**)*, and Aspergillus parasiticus* (**c**) pathogens in liquid culture in the presence of different concentrations of purified 5-rosanol, epoxyrosanol, and trihydroxyrosanol (THR). Assays were performed over a 48-hour time course in a defined RPMI-1640 MOPS buffer solution using a microtiter plate assay. Error bars represent standard error of the mean (n=6). Different letters (a-c) represent statistically significant differences in measurements at the 48 h time point; statistical analysis was performed using an ANOVA (p < 0.05), followed by Tukey’s Honest Significance Difference (HSD) to identify significantly different treatment pairs.

### Rosalexins are stress-inducible alongside other specialized maize diterpenoid pathways

To investigate associations of rosalexins with pathogen responses *in planta*, available maize genotypes featuring ZmTPS42/KSL1 genes demonstrated or predicted to be functional (Oh7B, Ki3, CML103, CML277) or containing non-functional *ZmTPS42/KSL1* genes (Oh43, B73, and B75) were selected for plant pathogen stress studies (**Fig. 2c**). Since rosalexins are significantly more abundant in stems as compared to roots (**Fig. 5**), analyses focused on stem tissue. Here, longitudinal stem incisions between nodes 3 and 4 of 62-day old plants were infected with either *F. verticillioides* or water (control). One week post infection, visual inspection showed that of lesion sizes caused by fungal infection varied across the tested maize genotypes with no distinguishable trends between lines featuring functional *vs* non-functional *ZmTPS42/KSL1*, respectively (**Fig. 7a**). Parallel transcriptome analysis of stem tissue showed that *ZmTPS42/KSL1* transcript abundance was lower as compared to *ZmTPS43/KSL2* and *ZmTPS45/KSL4*, which are the core genes controlling the committed kauralexin and dolabralexin pathway nodes, respectively. Shared pathway genes, including *ZmTPS38/CPS2/ZmAN2* and *ZmCYP71Z18* showed relatively higher gene expression than *ZmTPS42/KSL1* (**Fig. 7b; Supplementary Table S1**). In contrast, *ZmCYP71Z16*, which is only involved in the dolabralexin pathway, showed low expression in stem tissue, and had lower expression than *ZmTPS45/KSL1*. However, *ZmTPS42/KSL1* gene expression was increased in response to *F. verticillioides* in lines featuring functional genes as well as those lines carrying non-functional genes although to a somewhat lesser extent. Consistent with the only distant localization of *ZmTPS42/KSL1* and *ZmTPS40/CPS4* on chromosome 4, the two diTPS show clearly different expression patterns with the *ZmTPS40/CPS4* transcript being more abundant in root tissue without any clear pattern of pathogen inducibility. Genes in the zealexin, kauralexin, and dolabralexin pathways exhibited higher expression in response to fungal infection than control plants for all lines in stem tissues (**Fig. 7b, Supplementary Table S1)**.

**Fig. 7.**
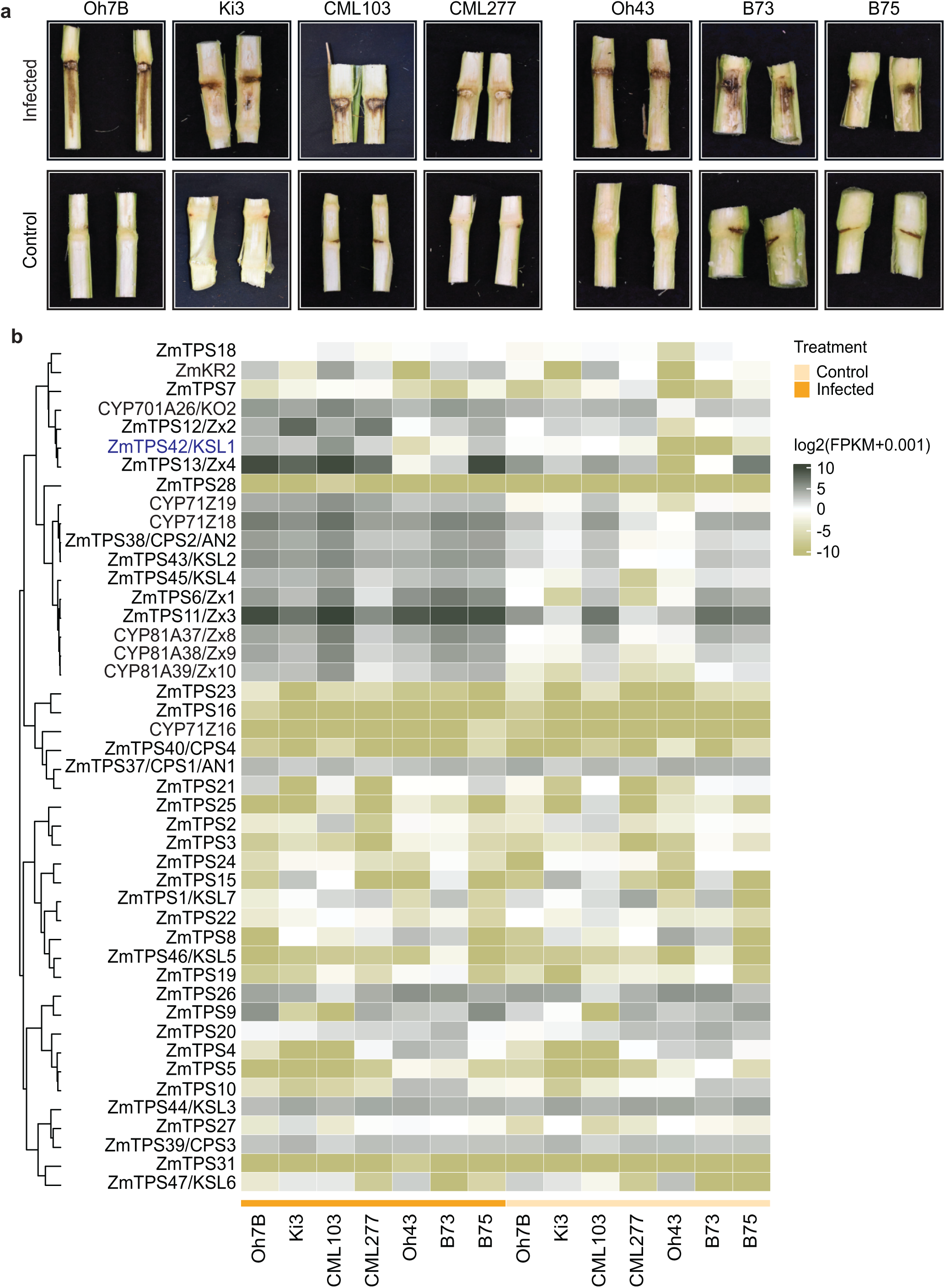
Rosalexins are pathogen-induced alongside other defense terpene pathways. a, Representative images of maize stems infected with *F. verticillioides* (infected) or water (control) using the lines Oh7B, Ki3, CML103, and CML277 (featuring a functional ZmTPS42/KSL1 gene) and Oh43, B73, and B75 (containing non-functional ZmTPS42/KSL1). **b,** heat map depicting gene expression (log2[FPKM+0.001]) of known maize terpene synthases, P450 genes, and kauralexin reductase 2 (KR2) in response to infection with *F. verticillioides* or mock treated controls (n=3).

## Discussion

Plants deploy species-specific networks of specialized metabolites to mediate dynamic interorganismal interactions and ecological adaptation. In light of rising environmental pressures that threaten crop health and production, understanding the metabolic and regulatory circuitry that contributes to innate immunity is pivotal for improving disease control and resistance trait breeding ^6,23,40^. Species-specific blends of diterpenoids with potent antimicrobial, insecticidal, and allelopathic functions that directly impact plant protection and vigor exist in major grain crops, including rice, maize, wheat, and barley ^5,9,41^. This study highlights the capacity of integrated multi-omics approaches and combinatorial enzyme biochemical tools to uncover hidden specialized metabolic pathways even in well-studied crops such as maize.

Prior maize research identified related yet distinct kauralexin and dolabralexin diterpenoids that serve as core components in the protection against fungal pathogens, insect pests, and drought stress ^16,17,19,21^. These diterpenoids arose through the functional expansion of the diTPS and P450 enzyme families via repeated events of often transposon-mediated tandem gene duplications and gene losses that occurred during the allopolyploidization and subsequent rediploidization events that shaped the maize genome ^42,43^. As a common mechanism in diterpenoid diversification, especially the duplication and divergence of GA-biosynthetic genes have contributed to diterpenoid chemical diversity ^12^. Indeed, such gene duplication events resulted in the evolution of two catalytically identical *ent*-CPP synthases, the GA-metabolic ZmTPS37/CPS1/AN1 and the strictly stress-elicited ZmTPS38/CPS2/AN2 ^25^. The distinct stress-responsive gene expression of *ZmTPS38/CPS2/AN2* enabled the dysregulation of GA and defensive diterpenoid metabolism in maize ^16,17^, and forms a metabolic hub for converting *ent*-CPP via different pathway modules. Here, ZmTPS43/KSL2 and ZmTPS45/KSL4 produce the committed *ent*-iso-kaurene and dolabradiene intermediates en route to kauralexins and dolabralexins, respectively ^16,17^ (**Fig. 1**). Phylogenetic relatedness to ZmTPS43/KSL2 and ZmTPS45/KSL4 suggests that ZmTPS42/KSL1 emerged as another gene duplication recruited from GA metabolism that functionally diverged toward specialized metabolism. The presence of functional and pseudogenized *ZmTPS42/KSL1* genes across different maize genotypes exemplifies the high genetic diversity in maize that shapes metabolic network architecture and chemical diversity (**Fig. 2**). The formation of rosalexins by the combined activity of ZmTPS38/CPS2/AN2, ZmTPS42/KSL1, and ZmCYP71Z18 adds yet another pathway branch to the portfolio of *ent*-CPP-derived diterpenoids produced in maize (**Fig. 1**). Due to their distinct product specificity, ZmTPS42/KSL1, ZmTPS43/KSL2, and ZmTPS45/KSL4 serve as core pathway branch nodes en route to kauralexins, dolabralexins, and rosalexins. Presence of ZmCYP71Z18 as a highly promiscuous P450 capable of functionally decorating each of the respective *ent*-iso-kaurene, dolabradiene, and 5-rosanol intermediates creates a modular pathway network for the efficient production of bioactive diterpenoid blends ^16,38^. The absence of significant co-regulation patterns (**Fig. 2 & 7**) and the low product abundance when pairing ZmTPS42/KSL1 with the 8,13-CPP synthase, ZmTPS40/CPS4, or the (+)-CPP synthase, ZmTPS39/CPS3 ^18^ (**Fig. 3**), support *ent*-CPP as the dominant ZmTPS42/KSL1 substrate. The formation of 5-rosanol by ZmTPS42/KSL1 reveals a previously unknown catalytic mechanism within the functional diversity of the class I diTPS family in plants. Based on known class I diTPS reactions and AI-assisted mechanistic studies we here propose that 5-rosanol formation proceeds by initial biosynthesis of a pimara-15-en-8-yl^+^ carbocation, followed by a series of 1,2-hydride and methyl migrations to yield the final rosa-15-en-5-yl^+^ intermediate, which is neutralized via water quenching (**Fig. 3**). As such, ZmTPS42/KSL1 expands the small group of known class I diTPSs that catalyze hydroxylation reactions during scaffold formation, including the *Salvia sclarea* scalerol synthase ^44,45^, *Isodon rubescens* nezukol synthase ^46^, and 9-hydroxy-pimarane synthases in switchgrass ^10^. Notably, small amounts of a predicted rosadiene scaffold suggest that the more prototypical deprotonation of the carbocation does occur during ZmTPS42/KSL1 catalysis (**Fig. 3 & 4**). Moreover, considering that additional C-19 methyl migration from the rosa-15-en-5-yl^+^ intermediate would lead to the formation of a dolabrane scaffold ^37^, supports a related evolutionary path of ZmTPS42/KSL1 and ZmTPS45/KSL4 consistent with their close phylogenetic relationship (**Fig. 2**). To current knowledge, the biosynthesis of 5-rosanol is unique to maize, although rosane-type diterpenoids have been identified in a range of, for example, liverworts as well as Araucariaceae, Linaceae, and Euphorbiaceae species ^47–50^.

The dependence of rosalexin accumulation on the presence of a functional *ZmTPS42/KSL1* gene sequence across the tested maize lines supports the hypothesis that *ZmTPS42/KSL1* represents the core pathway node for the biosynthesis of rosalexins (**Fig. 5**). Strikingly, similar to trihydroxydolabrene in dolabralexin metabolism ^17^, epoxyrosanol can undergo spontaneous ring opening yielding the trihydroxy derivative, THR, as a major metabolite *in planta* (**Fig. 5**). As similarly observed for dolabralexins, epoxyrosanol - but to lesser extend 5-rosanol or the THR - show potent antibiotic activity against all tested maize pathogens *F. verticillioides*, *A. parasiticus*, and *R. microsporus* (**Fig. 6**), highlighting that the epoxide moiety is critical for bioactivity in both dolabralexins and rosalexins. Notably, this finding aligns with the often high reactivity and antimicrobial potential of chemoenzymatically generated terpenoid epoxides ^51^. In maize, the functional relevance of these trihydroxy derivatives of epoxydolabrene and epoxyrosanol remains to be uncovered, given their limited antimicrobial activity but substantial abundance *in planta*. Despite the pathogen-elicited accumulation of rosalexins in stem tissue and the potent antimicrobial activity of epoxyrosanol *in vitro* (**Figs. 5 & 6**), the absence of a distinctive resistance phenotype in rosalexin-producing *vs* non-functional lines upon *F. verticillioides* infection suggest that rosalexins play a distinct context-dependent or lesser role in the dynamic network of maize chemical defense layers (**Fig. 7**). For example, consistent *ZmTPS42/KSL1* transcript accumulation patterns in aerial tissues have been observed in response to infection by other foliar pathogens such as *Cochliobolus heterostrophus* and *Colletotrichum graminicola* ^16,52^.

The discovery of the rosalexin pathway completes a triad of bioactive diterpenoid groups that evolved from ancestral GA metabolism in maize and predictably contribute to above- (kauralexins, rosalexins) and below-ground (dolabralexins) chemical defenses and other environmental interactions. Knowledge of these enzyme functions and their role in the modular maize diterpenoid-metabolic network forms the foundation to comprehensively understand what drives the substantial variation in the abundance, composition, and stress-inducibility of specialized diterpenoids across genetically diverse maize lines and commercial hybrids with direct impact on stress resilience ^17,19^. Ultimately, identifying maize lines with diterpenoid profiles promoting stress resilience provide resources for optimizing resistance traits.

## Material and Methods

### Plant and fungal material

Seeds of the tested maize genotypes were obtained from the Maize Genetic COOP Stock Center (https://maizecoopsc.org/) or the USDA National Plant Germplasm System (https://www.grin-global.org/) and cultivated as outlined below. *Fusarium verticillioides* GL1093 was kindly provided by Dr. Eric Schmelz, and isolates of *A. parasiticus* (NRRL #6433) and *R. microsporus* (NRRL #5558) were obtained from the Northern Regional Research Laboratory (NRRL). Pathogens were grown on potato dextrose agar for 3 days before the quantification and use of spores in bioactivity assays ^53,54^.

### Plant cultivation

For *in planta* detection of rosalexins (**Fig. 5**), plants were greenhouse grown at the University of California-San Diego Biology Field Station as previously described ^55^. In brief, maize seeds were sterilized using 70% EtOH followed by a 15% bleach solution before being washed with MilliQ water. Seeds were then germinated in MetroMix 200 (Sun Gro Horticulture Distribution) soil supplemented with 14-14-14 Osmocote (Scotts Miracle-Gro) and plants were cultivated with a 12-h photoperiod with a minimum of 300 μmol m^-2^s^-1^ of photosynthetically active radiation supplied by supplemental lighting, 70% relative humidity, and a day/night temperature cycle of 28°C/24°C. For rosalexin pathway profiling of selected maize lines (**Fig. 7**), plants were greenhouse grown at the University of California-Davis as described above with a few modifications. Seeds were germinated in an autoclaved soil mixture of half BM7 35% Bark HP and half Turface Athletics MVP and supplemented with nutrient water (Ca(NO_3_)_2_, KNO_3_, Fe-EDTA, Zn-EDTA, Cu-EDTA, Na_2_MoO_4_, KH_2_PO_4_, MgSO_4_, and K_2_SO_4_).

For enzyme functional assays, *N. benthamiana* was grown from seed in Sunshine Mix (SUN GRO) and supplemented with nutrient water (Ca(NO_3_)_2_, KNO_3_, Fe-EDTA, Zn-EDTA, Cu-EDTA, Na_2_MoO_4_, KH_2_PO_4_, MgSO_4_, and K_2_SO_4_) in controlled environment growth chambers under a photoperiod of 12/12 h, 60% relative humidity, 100 μM m^−2^s^−1^ light intensity, and a day/night temperature cycle of 26/25°C.

### Fungal pathogen infection of maize tissues

For stem inoculations with *F. verticillioides*, 8-11 cm longitudinal incisions were made between nodes 3 and 4 using a surgical scalpel and treated with 1 mL of fungal spore solution (1×10^7^ mL^−1^) in MilliQ water or with water only as a control. For stem lesion imaging, a BD PrecisionGlide Needle (18G) was used to create a 1.2 mm diameter hole under node 3 and inoculated with 1 mL of fungal spore solution (1×10^7^ mL^−1^) in MilliQ water. Wound sites were wrapped with clear packing tape to prevent desiccation. For root elicitation assays, soil was gently brushed aside to reveal nodal roots, which were punctured at 1 cm intervals using a BD PrecisionGlide needle (27G1/2) and inoculated with 50 µL fungal spore solution (1×10^7^ mL^−1^) in MilliQ water or water only (control) at each wound site. Plants were additionally watered with 600 mL of fungal spore solution (1×10^7^ mL^−1^) in MilliQ water or water (control) to increase root infection. Plants (n=3-8 per genotype) were harvested 7-9 days post infection over the course of 3 days in a random order to account for variation that may have occurred over the collection period. Representative root sections were collected and washed gently with deionized water. Stem tissue was collected using sterile garden sheers and a Zyliss 3.25 inch paring knife. Lesion images were taken immediately after harvest. Stem tissue from both wounding methods were pooled together for each plant for use in metabolomics and transcriptomics studies. All tissue samples were immediately frozen in liquid N_2_ and stored at -80℃ for further analysis.

### Transcriptome sequencing and analysis

For transcriptome analysis, root and stem tissue (n=3) from control and fungal infected plants of maize genotypes selected for the pathogen elicitation assays described above was used. RNA was extracted using the Quick-RNA Plant Miniprep kit (Zymo Research). Gene expression analysis of the RNA-seq data was performed by Novogene. In brief, RNA sequencing libraries were prepared and sequenced using the Illumina NovaSeq X-Plus Sequencing platform with a 2x150 paired-end read configuration. Raw sequencing data were generated as FASTQ files. A total of 84 libraries were sequenced, yielding raw read counts ranging from 39,892,816 to 76,617,254 reads per sample. The obtained data were processed to remove low-quality sequences and adapter contamination using the Novogene QC pipeline. After filtering, the number of retained reads ranged from 38,794,843 to 67,827,953 reads per sample. Filtered reads from each library were aligned to the maize reference genome (Zm-B73-REFERENCE-NAM-5.0) using HISAT2 version 2.2.1 ^56^. FeatureCounts (2.06) ^57^ was used to count the number of reads mapped to each gene. The resulting averaged Fragments per Kilobase of transcript Million (FPKM) mapped reads were log2[FPKM+0.001] transformed and analyzed in RStudio using the ComplexHeatmap package to generate the gene expression heat map.

### *In vivo* transient co-expression assays in *Nicotiana benthamiana*

Full-length, codon optimized genes of the focal diTPSs and P450s were obtained by DNA synthesis (Twist Bioscience) with financial support by the DOE Joint Genome Institute (JGI) DNA Synthesis Science Program (grant no. 510448) and inserted into the binary pEAQ-HT vector system ^58^. For co-expression in *N. benthamiana*, constructs were transformed into *A. tumefaciens* strain GV3101 and cells were grown at 28°C for 48 hours in LB media supplemented with 10 µL mL^-1^ of gentamicin and rifampicin and 50 µL mL^-1^ of kanamycin. Cells were then adjusted to a final optical density (OD_600_) of 0.5 in 10 mM MES buffer (pH 5.6) with 10 mM MgCl_2_ and 150 µM acetosyringone (Acros Organics). For co-expression of the chosen diTPS and P450 combinations the respective Agrobacterium cultures were combined in equal volumes including the silencing suppressor strain p19 ^59^ and infiltrated into the abaxial side of the leaves of 5-week-old *N. benthamiana* plants using a needless syringe. Five days post-infiltration, metabolite extraction from transfected plants was performed via a 15-minute sonication using 5 mL 85:15 (v/v) hexane:ethyl acetate (Fisher Scientific) and analyzed via GC-MS or LC-MS/MS as described below.

### GC-MS analysis of enzyme products

GC-MS analysis of enzyme products was conducted on an Agilent 7890B GC interfaced with a 5977 Extractor XL MS Detector at 70 eV and 1.2 mL min^−1^ He flow, using an Agilent DB-XLB column (30 m, 250 µm i.d., 0.25 µm film) and the following GC parameters: 50°C for 3 min, 15°C min^−1^ to 300°C, hold for 3 min with pulsed splitless injection at 250°C. MS data from 40– 400 mass-to-charge ratio (*m*/*z*) were collected after a 13-min solvent delay. Metabolite quantification (n=4) was based on normalization to the internal standard 1-eicosene (Sigma-Aldrich), tissue dry weight, and OD_600_ at the time of induction of protein expression.

### Metabolite profiling of maize tissues via LC-MS/MS

To quantify plant metabolites, 100 mg of maize tissue was collected from the fungal elicitation experiment (see above). Fresh tissue was lyophilized for 48 h and then ground into a fine powder using a ball Retsch mill. Metabolites were extracted using 80% Optima LC-MS grade MeOH (Fisher Scientific) containing 0.5 µM Telmisartan (Sigma-Aldrich) as internal standard for quantification. Liquid chromatography was performed on a Thermo Scientific Vanquish UHPLC with an Acquity UPLC BEH C18 column (2.1 × 100 mm, 1.7 μm particle size; Waters) using Optima LC-MS grade water with 0.1% formic acid (v/v) as eluent A and Optima acetonitrile with 0.1% formic acid (v/v) as eluent B. The chromatography method used a flow rate of 0.6 mL min^-1^: -1 to 0 min, 1% B, 0 to 0.5 min, 50% B, 0.5 to 7.5 min, 90% B, 7.5 to 9 min, 1% B, 9 to 9 min 1% B, 9 to 10 min 1% B with the column compartment set to 40°C. Mass spectrometry was performed using a Thermo Scientific Q-Exactive equipped with an electrospray ionization source that was run in positive ionization mode (spray voltage, 3.50 |kV|; capillary temperature, 300°C; aux. gas heater, 350°C; sheath gas flow rate, 45; aux. gas flow rate, 10; sweep gas flow rate, 3). The MSDial software program ^60^ was used to generate and analyze the LC-MS/MS generated data. To quantify metabolites, peak area was autointegrated and extracted from Thermo Scientific Free Style and all calculations were performed in Excel (**Supplementary Table S6**).

### *In vitro* antifungal assays

Fungal growth inhibition assays were performed following the Clinical and Laboratory Standards Institute M38-A2 guidelines as described previously ^20^. Fungal isolates were cultured on potato dextrose agar for three days at room temperature before the quantification and use of spores. Spore densities were estimated using a hemocytometer and initial fungal inoculum was diluted with RPMI-1640 MOPS buffer (pH 7) to 2.5 × 10^4^ conidia mL^-1^. Fungal growth at 30°C was monitored using a Infinite M Nano microplate absorbance reader (Tecan) with a 96-well microtiter plate-based method. Each well contained 200 µL of initial fungal inoculum with 2.5 µL of pure DMSO or RXs in DMSO at concentrations of 10, 25, and 50 µg mL^-1^. Negative and positive controls consisted of 200 µL of RPMI-1640 MOPS or initial fungal inoculum, respectively, each containing 2.5 µL of pure DMSO. Changes in OD_600_ were determined using averages (n=6) of OD_600_ values measured hourly over 48 hours. Fungal culture OD_600_ values were normalized first to the starting OD_600_ at time point 0, then to the OD_600_ of the negative control at each corresponding time point. Data were assessed via Analysis of Variance (ANOVA) followed by a Tukey Honest Significant Difference (HSD) statistical analysis. Area under the curve (AUC), approximated under the Trapezoidal Rule, was used to calculate fold change in growth for the statistically significant concentrations relative to the positive control. Uncertainty was estimated by Monte Carlo simulations [CI 95%, n = 5000] to propagate replicate variation into AUC comparisons.

### Chemical synthesis and purification of trihydroxyrosanol from epoxyrosanol

Trihydroxyrosanol (THR) was generated by acid-catalyzed epoxide opening of epoxyrosanol. Purified epoxyrosanol (2 mg) was dissolved in 4 mL of 60% (v/v) acetone and vortexed thoroughly. The reaction was initiated by adding 220 µL of 0.5 M H₂SO₄ and incubating the mixture at room temperature for approximately 12 h. The reaction was quenched by the dropwise addition of concentrated NaHCO₃ until effervescence ceased, followed by the addition of 4 mL saturated NaCl brine. The product was extracted three times with 1 mL of 9:1 (v/v) ethyl acetate:methanol. Combined organic layers were washed with 600 µL brine, dried over anhydrous Na₂SO₄, and transferred to a clean vial. Conversion and purity were verified by LC-MS/MS. Absolute purification was performed using a semi-preparative Agilent 1100 Series HPLC system equipped with an Agilent Eclipse XDB-C18 column (5 µm, 4.6 × 150 mm). A gradient elution was performed with solvent A (Milli-Q water) and solvent B (HPLC-grade acetonitrile) at a flow rate of 1 mL min⁻¹. The gradient profile was 0 min 20% B to 30 min 40% B. The purified THR fraction was collected and analyzed by NMR in MeOH-d₄ as described below.

### Experimental NMR analysis

For NMR analysis, ≥1 mg of diterpenoid products was enzymatically produced via transient expression in *N. benthamiana* as outlined above. The reaction products were extracted in 85:15 (v/v) hexane:ethyl acetate and purified through iterative separation on a silica matrix with a hexane/ethyl acetate gradient as the mobile phase. Following separation via a 500 mL silica matrix column, products were further purified via semi-preparative Agilent 1100 Series HPLC equipped with an Agilent Eclipse XDB-C18 5 µm 4.6x150 mm column using a gradient elution with eluent A 100% MilliQ water and eluent B HPLC-grade acetonitrile. Products were eluted under the following conditions: Epoxyrosanol: 0 min 60% B to 30 min 90% B; 5-rosanol: 0 min 80% B to 30 min 100% B; epoxyrosaene: 0 min 50% B to 30 min 100% B with 1 mL fractions collected over 30 min at a flow rate of 1 mL min^-1^. Purified products were dissolved in deuterated chloroform-d (CDCl_3_; Sigma-Aldrich) containing 0.03% (v/v) tetramethylsilane (TMS). The 1D (1H, 13C, NOE) and 2D (HSQC, COSY, HMBC, H2BC and NOESY) spectra were acquired on a Bruker Avance III 800 MHz spectrometer equipped with a 1H-13C / 15N / 2H Bruker CPTCI Cryoprobe. NMR spectra were analyzed using Bruker TOPSPIN 3.2.

### Computational stereochemical analysis of 5-rosanol

The target diastereomer was predicted by comparing experimental proton and carbon NMR chemical shift values to calculated ones for the 16 possible structures ^61^. For each diastereomer, the method begins with a conformational search. Low energy conformations were found using xTB-CREST, then optimized using Gaussian 16 with a restricted B3LYP-D3(0)/6-31+G(d,p) level of theory and implicit solvation model (IEFPCM) for chloroform ^62–67^. Conformations within 3 kcal/mol of the lowest energy conformers were used to calculate computational NMR shifts using the GIAO method at the mPW1PW91/6-311+G(2d,p), and chloroform as the solvent ^68,69^. Both proton and carbon isotropic shielding values were computed. These computed isotropic shielding values were converted using CHESHIRE to predicted NMR shifts which were weighted based on their calculated free energy in the optimization calculation and averaged using the referenced code^61^. The predicted NMR shifts for each of the diastereomers were compared to the experimental NMR shifts using DP4+ to determine which diastereomer’s predicted shifts were most similar to the experimental ^70^. This method resulted in isomer 11 having the closest predicted shifts to the experimental with a computed probability of 100%.

### Computational analysis of the *Zm*TPS42/KSL1 reaction mechanism

The free energy profile for the formation of 5-rosanol was generated using RxnNet ^71^, an AI-assisted automated reaction mechanism discovery platform. RxnNet requires only the SMILES corresponding to the initial and final carbocations as the input. It automatically identifies the reaction cascade between the reactant and product and constructs the corresponding free energy diagram. RxnNet generates the minimum energy path (MEP) for each step of the reaction in a stepwise manner, and generates near-attack conformers (NAC) ^32,33^ for intermediates prior to performing the actual chemical step. For all chemical and conformational steps, an initial MEP guess is generated with the help of geodesic interpolation, followed by refinements using climbing image nudged elastic band (NEB-CI) calculations. A NEB-CI is initially performed at the semi-empirical quantum chemistry (QC) level of theory using GFN2-XTB ^32^. The final MEP is then obtained from NEB-CI using M06-2X/def2-SVP level of theory ^72–74^, and based on this MEP RxnNet identifies stationary points. RxnNet employs ORCA 6.0.1 ^75,76^ to perform all QC calculations. Readers can refer to reference ^71^ for further details on RxnNet.

### Mutual rank and Pearson correlation coefficient calculation of genes

RNA-seq datasets of infected and control (n=3) stem tissue from maize lines Oh7B and Ki3 were used for analyzing Mutual Rank (MR) and Pearson correlation coefficients (PCC). Raw FPKM values were log2[FPKM+0.001] transformed and zero variance genes were removed prior to analysis. *ZmTPS38/CPS2/AN2* and *ZmTPS42/KSL1* were used as bait genes and tested against *kauralexin reductase 2* (*KR2*) as well as TPS and CYP genes involved in specialized metabolism and GA biosynthesis. PCC values were calculated using pairwise complete observations. To assess correlation strength and specificity, a MR analysis was performed for each gene pair. PCC values were generated for each gene in the gene pair against all expressed genes in the dataset, and the MR value was calculated from the geometric mean between these two ranks. Gene pair strength was visualized using heat maps. Strong correlations were determined by a MR<200 and a PCC >0.7 and pairs that met the cutoff were shaded in orange using -log10(MR) whereas genes below the cut off were characterized as weak correlation and shaded in grey using -1og10(MR). Stars denote particularly strong correlations with a PCC>0.8 and MR <50.

### Sequence and Phylogenetic Analysis

Sequence alignments of diTPS genes were generated using the CLCBio Workbench (Qiagen) with default parameters. A maximum-likelihood phylogenetic tree was generated using PhyML ^77^ with four rate substitution categories, LG substitution model, BIONJ starting tree, and 500 bootstrap repetitions.

## Supporting information

Supplementary Fig. S1: Protein sequence alignment of ZMTPS42/KSL1 derived from selected maize lines.

Supplementary Fig. S2: Mass spectra of products from the pairwise reaction of ZTPS42/KSL1 with different maize class II diterpene synthases.

Supplementary Fig. S3: NMR structural elucidation of compound 1, 5-rosanol.

Supplementary Fig. S4: MEP plots of the chemical steps in the ZmTPS42/KSL1-catalyzed mechanism, as obtained from RxnNet.

Supplementary Fig. S5: NMR structural elucidation of compound 13, epoxyrosanol.

Supplementary Fig. S6: NMR structural elucidation of compound 14, epoxyrosaene.

Supplementary Fig. S7: Functional analysis of maize P450 enzymes for 5-rosanol conversion.

Supplementary Fig. S8: NMR structural elucidation of compound 15, trihydroxyrosanol (THR).

Supplementary Table S1: FPKM values associated with gene expression analysis of diterpenoid-metabolic pathways.

Supplementary Table S2: Stereochemistry of tested computational NMR structures for 5-rosanol

Supplementary Table S3: Computational NMR chemical shifts for 5-rosanol.

Supplementary Table S4: MEPs associated with the chemical steps in the ZmTPS42/KSL1-catalyzed mechanism, as obtained from RxnNet.

Supplementary Table S5: MS feature area alignment.

Supplementary Table S6: LC-MS/MS quantification calculations.

Supplementary Table S7: Raw reads from Tecan analysis of fungal bioactivity assays

## Data availability

The RNA-sequencing data reported in this study were submitted to the Sequence Read Archive (SRA), accession no. SUB16112077. Reported NMR data have been deposited at ioChem-Find: Home (iochem-bd.org). Computational NMR files have been deposited here: https://iochem-bd.bsc.es/browse/handle/100/492343.

## Supporting Information

Additional supporting information may be found online in the Supplementary Data section at the end of the article.

**Supplementary Fig. S1:** Protein sequence alignment of ZMTPS42/KSL1 derived from selected maize lines.

**Supplementary Fig. S2:** Mass spectra of products from the pairwise reaction of ZTPS42/KSL1 with different maize class II diterpene synthases.

**Supplementary Fig. S3:** NMR structural elucidation of compound 1, 5-rosanol.

**Supplementary Fig. S4:** MEP plots of the chemical steps in the ZmTPS42/KSL1-catalyzed mechanism, as obtained from RxnNet.

**Supplementary Fig. S5:** NMR structural elucidation of compound **13**, epoxyrosanol.

**Supplementary Fig. S6:** NMR structural elucidation of compound **14**, epoxyrosaene.

**Supplementary Fig. S7:** Functional analysis of maize P450 enzymes for 5-rosanol conversion.

**Supplementary Fig. S8:** NMR structural elucidation of compound **15**, trihydroxyrosanol (THR).

**Supplementary Table S1:** FPKM values associated with gene expression analysis of diterpenoid-metabolic pathways.

**Supplementary Table S2:** Stereochemistry of tested computational NMR structures for 5-rosanol

**Supplementary Table S3:** Computational NMR chemical shifts for 5-rosanol.

**Supplementary Table S4:** MEPs associated with the chemical steps in the ZmTPS42/KSL1-catalyzed mechanism, as obtained from RxnNet.

**Supplementary Table S5:** MS feature area alignment.

**Supplementary Table S6:** LC-MS/MS quantification calculations.

**Supplementary Table S7:** Raw reads from Tecan analysis of fungal bioactivity assays

## Acknowledgement and funding sources

We dedicate this work to the memory of Dr. Eric A. Schmelz (EAS.), whose mentorship, insight, and scientific vision contributed critically to this study. His passing is a great loss to the scientific community. We would like to thank Dr. Juan Carlos Caravez at the University of California, Santa Barbara for his expertise and guidance in synthesizing THR from epoxyrosanol for use in fungal bioactivity assays. Financial support for this work was provided by the National Science Foundation Plant Biotic Interaction (PBI) program (grant # 1758976 to EAS and PZ) and TRTech-PGR program (grant# 2312181 to PZ), as well as the United States-Israel Binational Agricultural Research and Development (BARD) Fund (grant # US-5682-24 to PZ and DTM. Additional financial support for this work by the US Department of Energy (DOE) Joint Genome Institute (JGI) DNA Synthesis Science Program (grant no. 510448, to PZ) is gratefully acknowledged. The gene synthesis work conducted by the JGI, a DOE Office of Science User Facility, is supported under contract no. DE-AC02-05CH11231.

## Notes

**Competing Interest Statement:** The authors declare no competing interests.

### Competing Interest Statement

The authors have declared no competing interest.

