## Supplementary Fig. S1: Protein sequence alignment of ZMTPS42/KSL1 derived from selected maize lines. for "Discovery of the rosalexin pathway expands the modular network of maize diterpenoid chemical defenses"

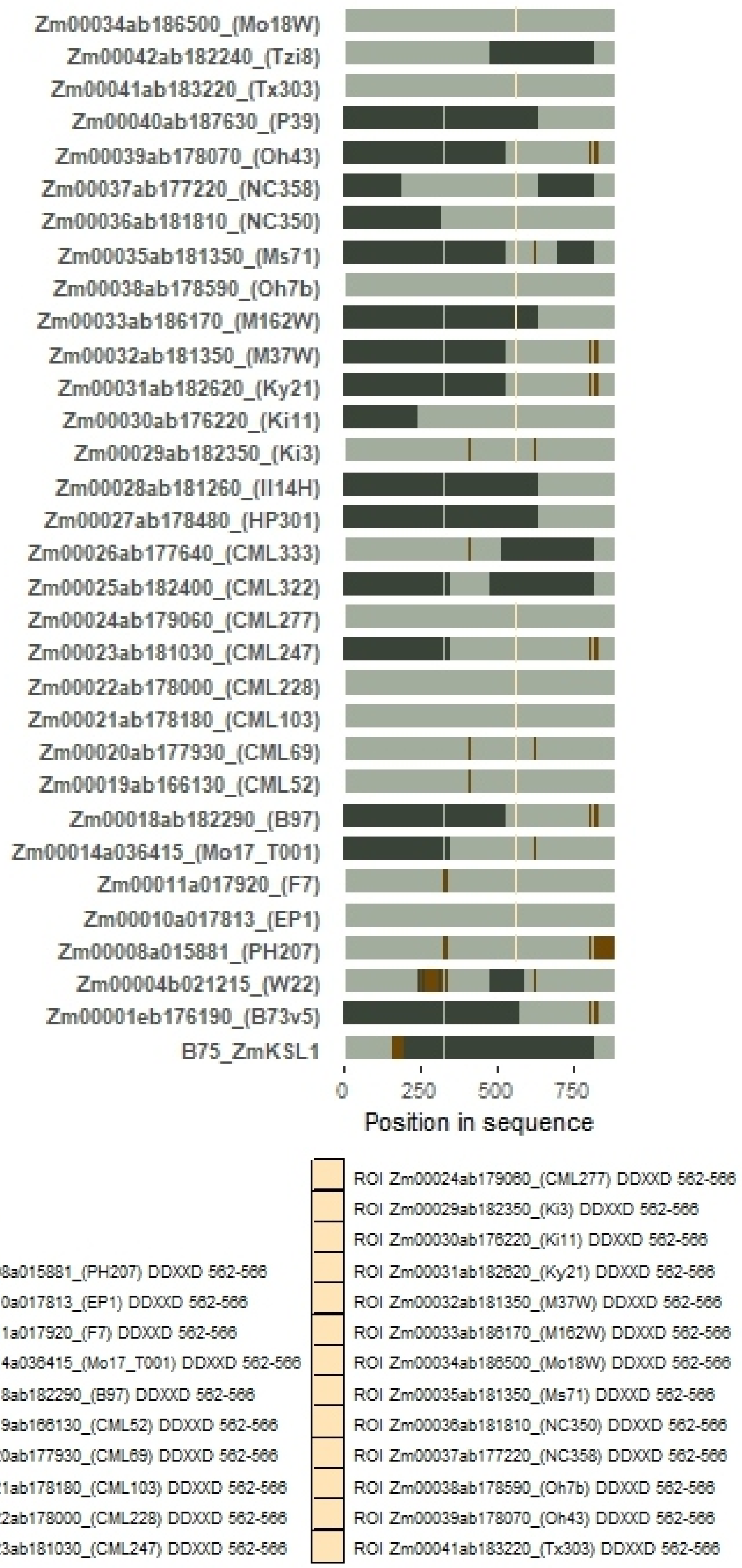


**Supplementary Fig. S1 | Protein sequence alignment of ZmTPS42/KSL1 sequences derived from selected maize lines.**
