## Supplementary Fig. S2: Mass spectra of products from the pairwise reaction of ZTPS42/KSL1 with different maize class II diterpene synthases. for "Discovery of the rosalexin pathway expands the modular network of maize diterpenoid chemical defenses"

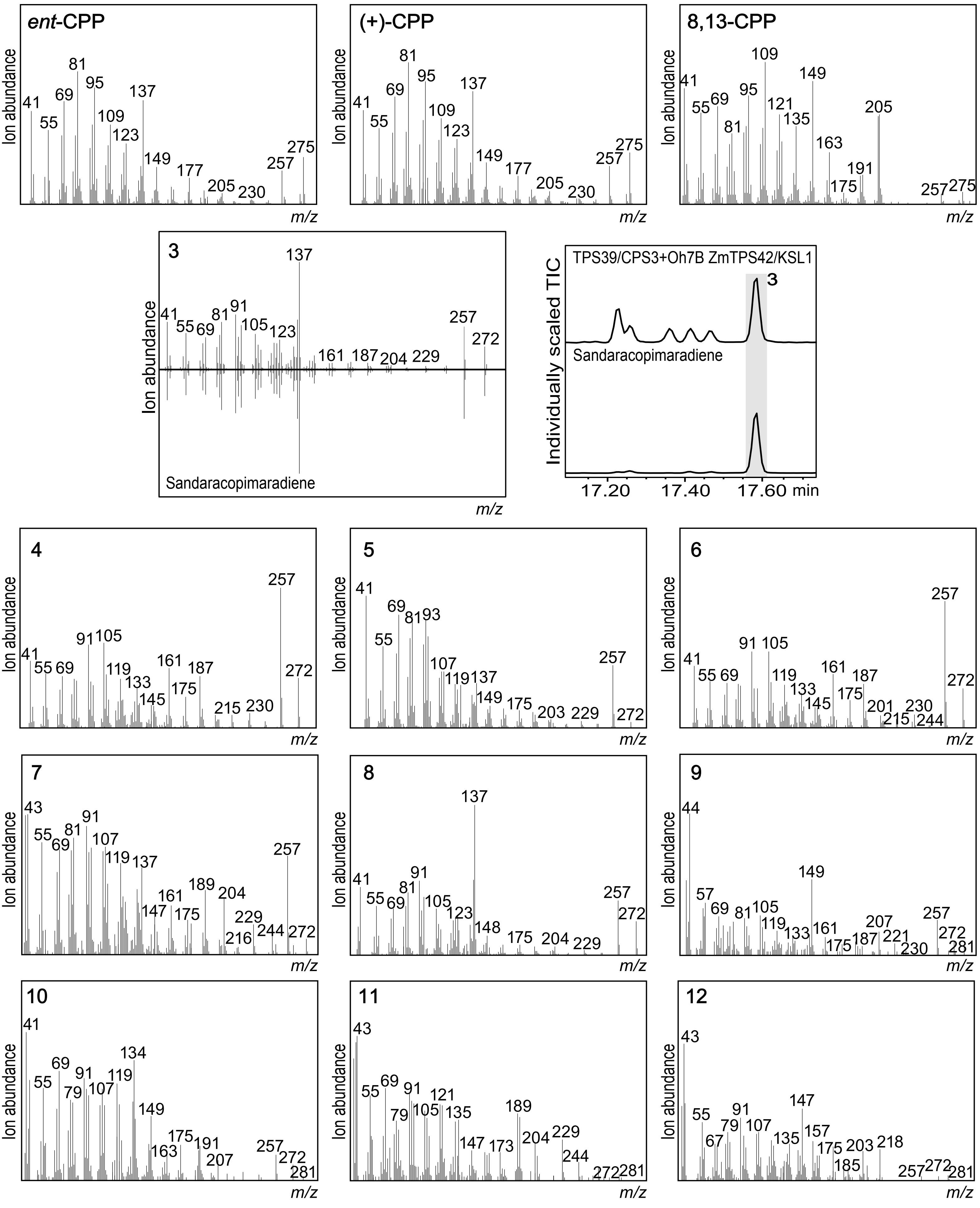


**Supplementary Figure S2 | Mass spectra of products from the pairwise reaction of ZmTPS42/KSL1 with different maize class II diterpene synthases.** Mass spectrum of Agrobacterium-mediated *Nicotiana benthamiana* co-expression assays of ZmTPS38/CPS2/AN2, ZmTPS39/CPS3, and ZmTPS40/CPS4 alone as well as in combination with ZmTPS42/KSL1. *Ent*-CPP is the major product of ZmTPS38/CPS2/AN2. (+)-CPP is the major product of ZmTPS39/CPS3 and compounds **4** and **5** are minor products. Compounds **3-8** were found in the ZmTPS39/CPS3 + ZmTPS42/KSL1 combination and compounds **3**, **6-8** are unique to ZmTPS39/CPS3 + ZmTPS42/KSL1 combination. Compound **3** was identified as sandaracopimaradine by using a standard. 8,13-CPP is the major product of ZmTPS40/CPS4 and compounds **9 - 12** are minor products.
