## Supplementary Fig. S3: NMR structural elucidation of compound 1, 5-rosanol. for "Discovery of the rosalexin pathway expands the modular network of maize diterpenoid chemical defenses"

**a**

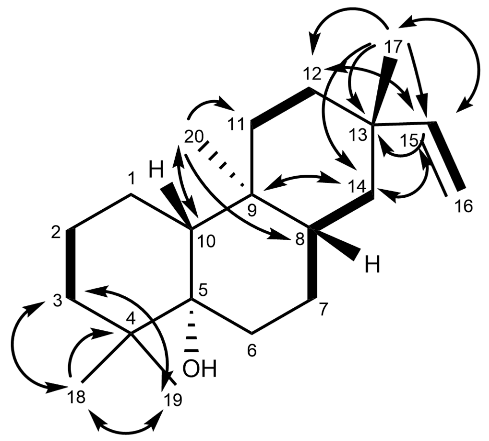


**b**

| Position | δC (ppm) | δH (ppm) |
|---|---|---|
| 1 a | | 1.626 |
| b | 21.183 | 1.358 |
| 2 a | | 1.589 |
| b | 22.094 | 1.477 |
| 3 a | | 1.665 |
| b | 36.906 | 1.121 |
| 4 | 38.843 | |
| 5 | 76.675 | |
| 6 a | | 1.722 |
| b | 32.623 | 1.572 |
| 7 a | | 1.524 |
| b | 25.07 | 1.121 |
| 8 | 41.669 | 1.2869 |
| 9 | 36.688 | |
| 10 | 48.63 | 1.338 |
| 11 a | | 1.585 |
| b | 35.083 | 1.111 |
| 12 a | | 1.578 |
| b | 31.95 | 1.204 |
| 13 | 36.361 | |
| 14 a | | 1.422 |
| b | 39.21 | 1.003 |
| 15 | 151.426 | 5.831 |
| 16 a | | 4.934 |
| b | 108.574 | 4.859 |
| 17 | 23.1 | 1.0507 |
| 18 | 24.42 | 0.884 |
| 19 | 24.01 | 1.0342 |
| 20 | 12.004 | 0.919 |

c

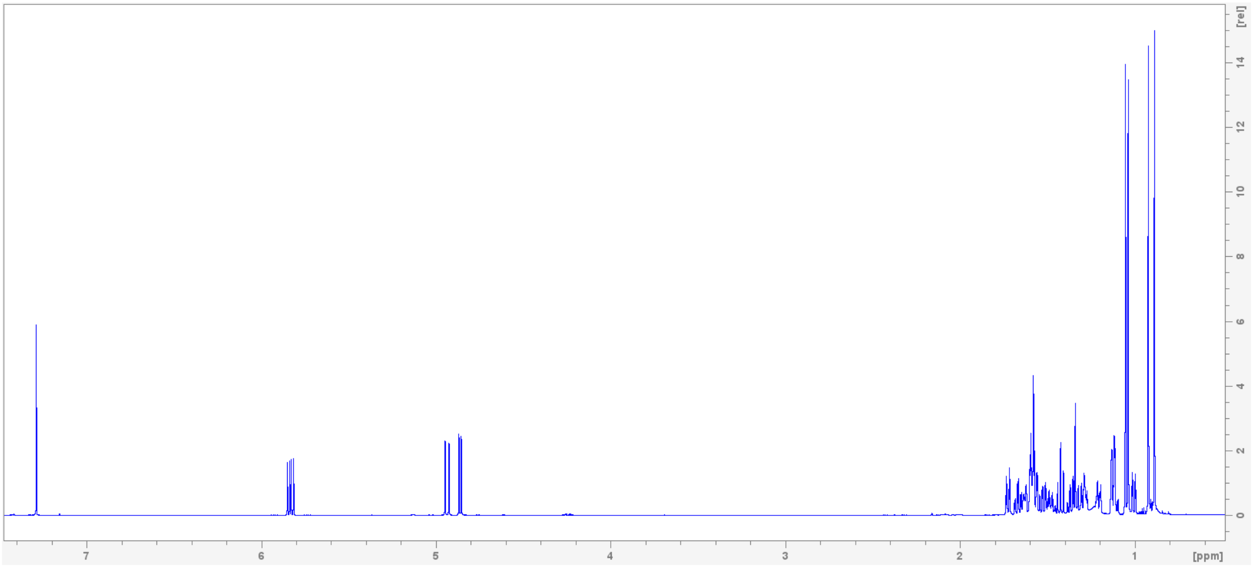
[rel]
14
12
10
8
6
4
2
0
7
6
5
4
3
2
1
[ppm]

d

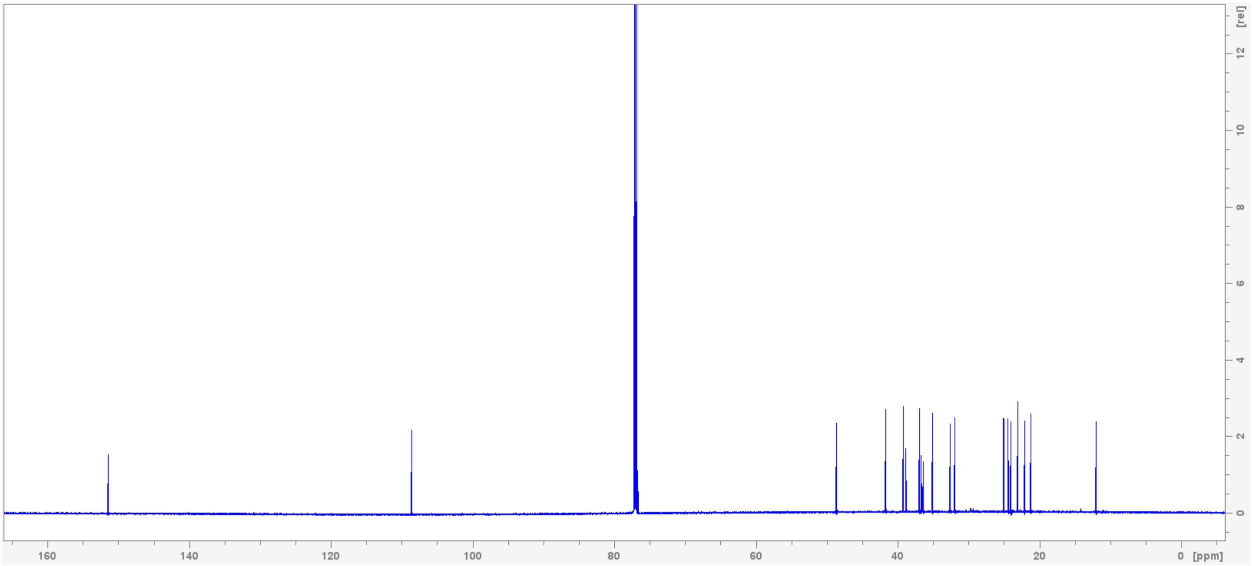
[rel]
12
10
8
6
4
2
0
160
140
120
100
80
60
40
20
0
[ppm]

e

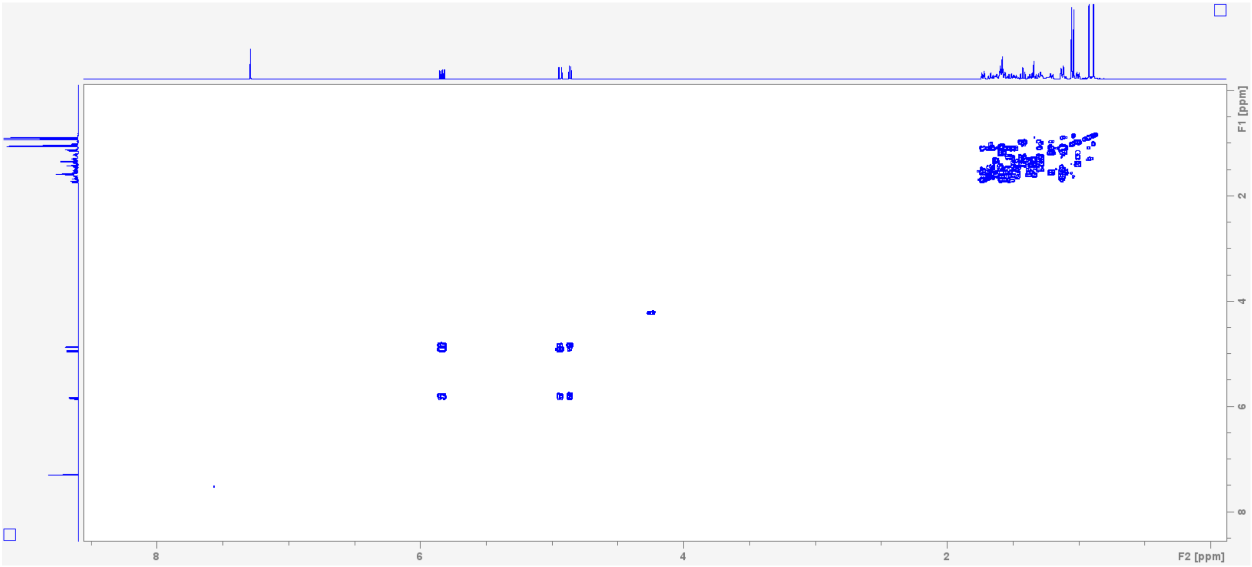
F1 [ppm]
2
4
6
8
8
6
4
2
F2 [ppm]

f

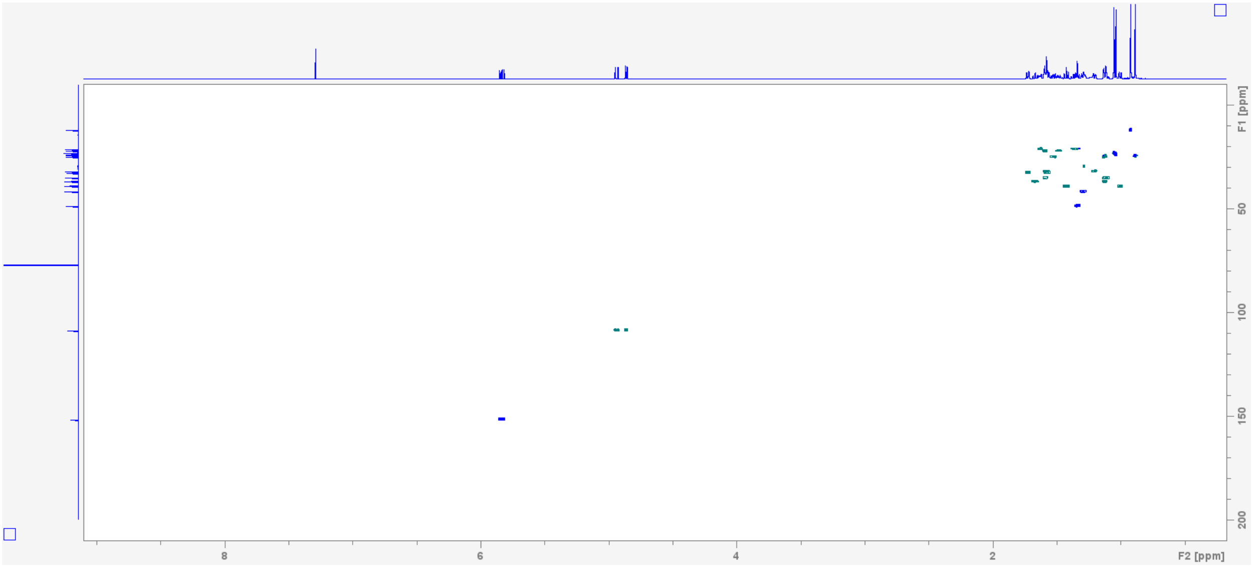


g

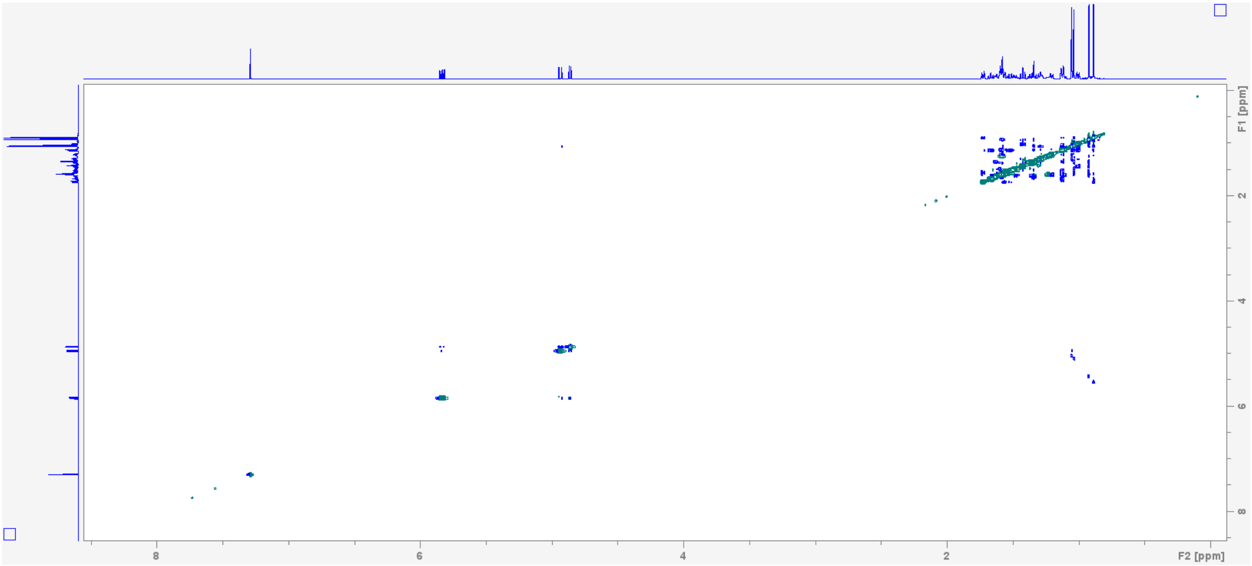


**Supplementary Figure S3.** NMR structural elucidation of compound **1**, 5-rosanol. **a,** chemical structure depicted with key structural elements solved by experimental NMR including COSY (bold) and HMBC (arrows). Other structural elements including stereochemistry were confirmed utilizing computation NMR. **b**, 5-rosanol $^{13}C$ and $^{1}H$ assignments and chemical shift table. 5-rosanol NMR spectra: **c**, $^{1}H$ NMR; **d**, $^{13}C$ NMR; **e**, HSQC; **f**, COSY; **g**, NOESY.
