## Supplementary Fig. S4: MEP plots of the chemical steps in the ZmTPS42/KSL1-catalyzed mechanism, as obtained from RxnNet. for "Discovery of the rosalexin pathway expands the modular network of maize diterpenoid chemical defenses"

**a**

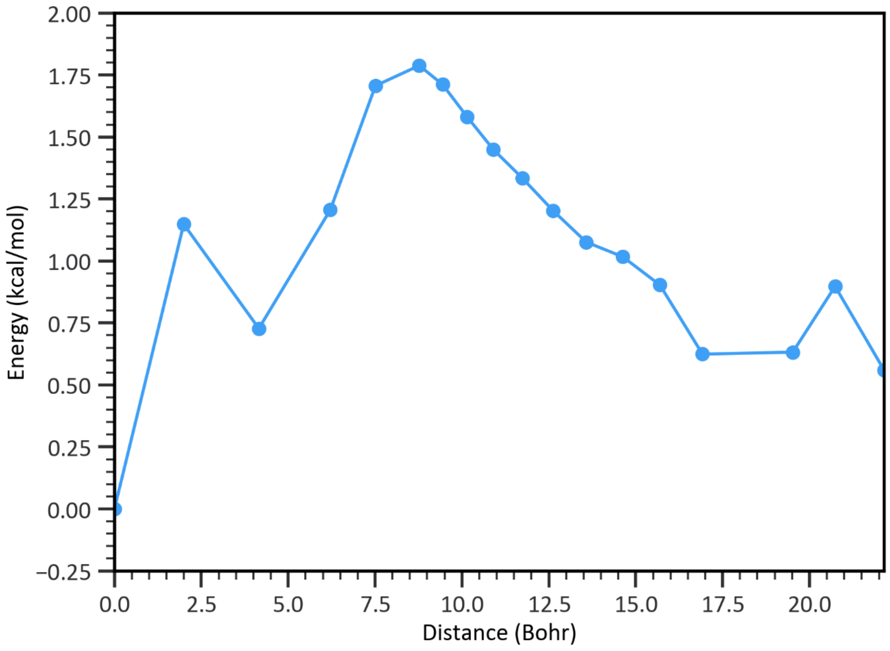
Energy (kcal/mol)
Distance (Bohr)
2.00
1.75
1.50
1.25
1.00
0.75
0.50
0.25
0.00
−0.25
0.0
2.5
5.0
7.5
10.0
12.5
15.0
17.5
20.0

**b**

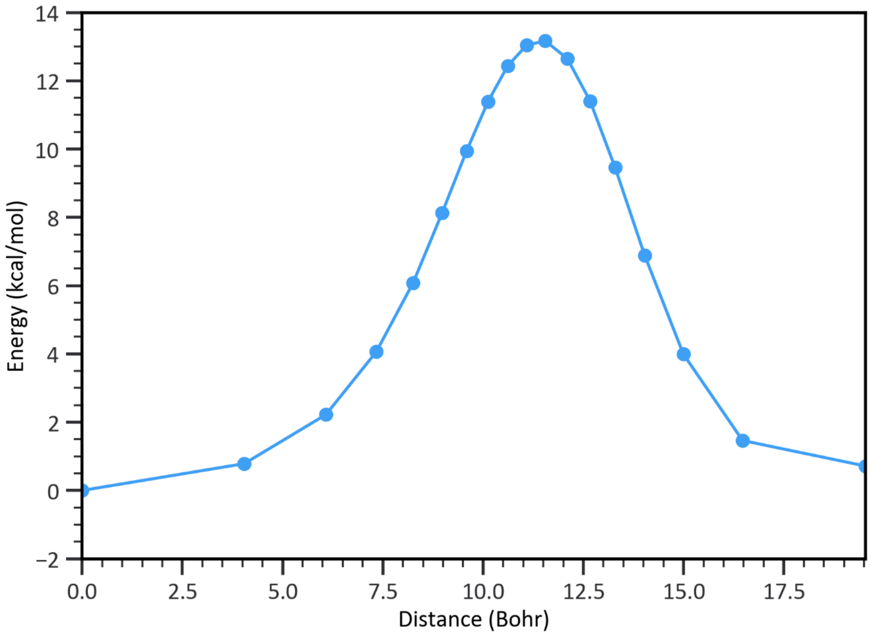
Energy (kcal/mol)
Distance (Bohr)
14
12
10
8
6
4
2
0
−2
0.0
2.5
5.0
7.5
10.0
12.5
15.0
17.5

c

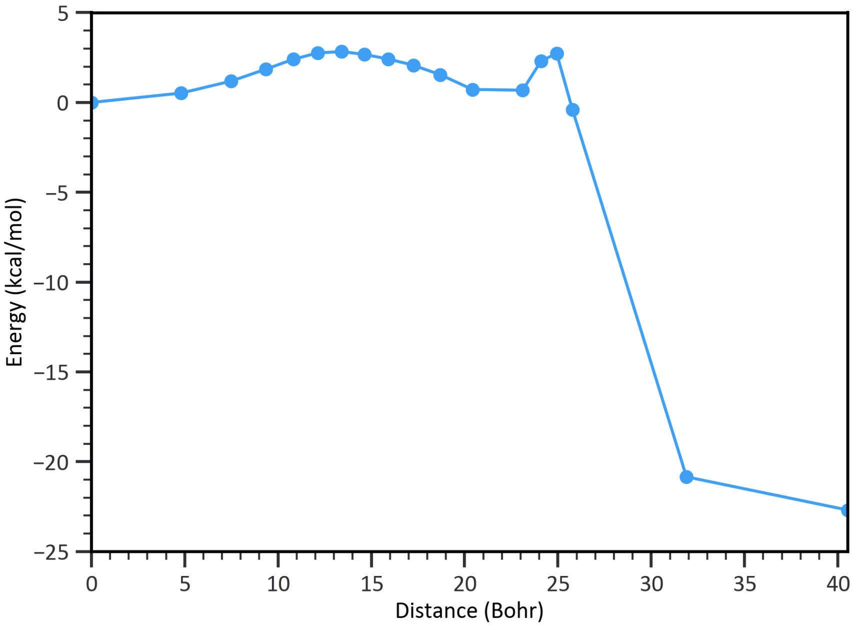

5
0
−5
−10
−15
−20
−25
Energy (kcal/mol)
0
5
10
15
20
25
30
35
40
Distance (Bohr)


d

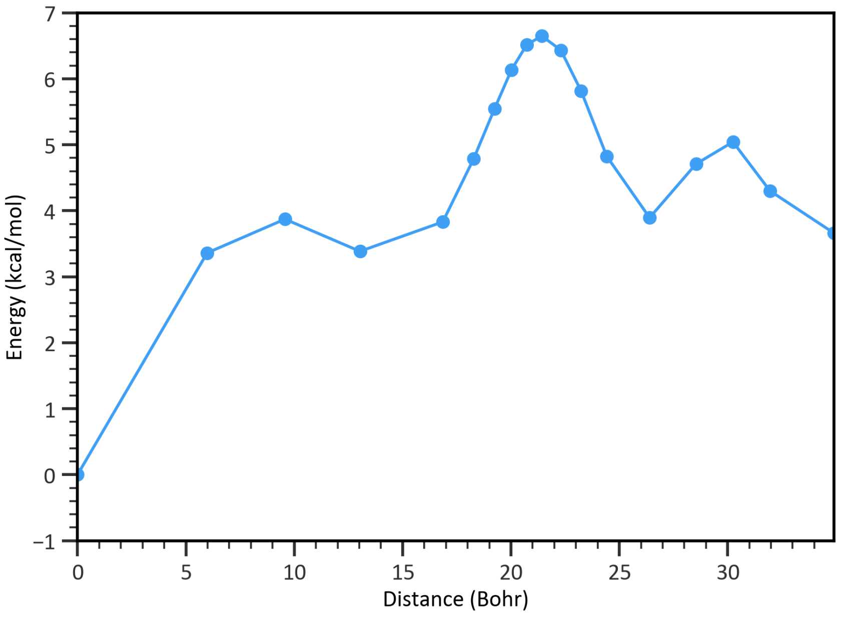

7
6
5
4
3
2
1
0
−1
Energy (kcal/mol)
0
5
10
15
20
25
30
Distance (Bohr)

**e**

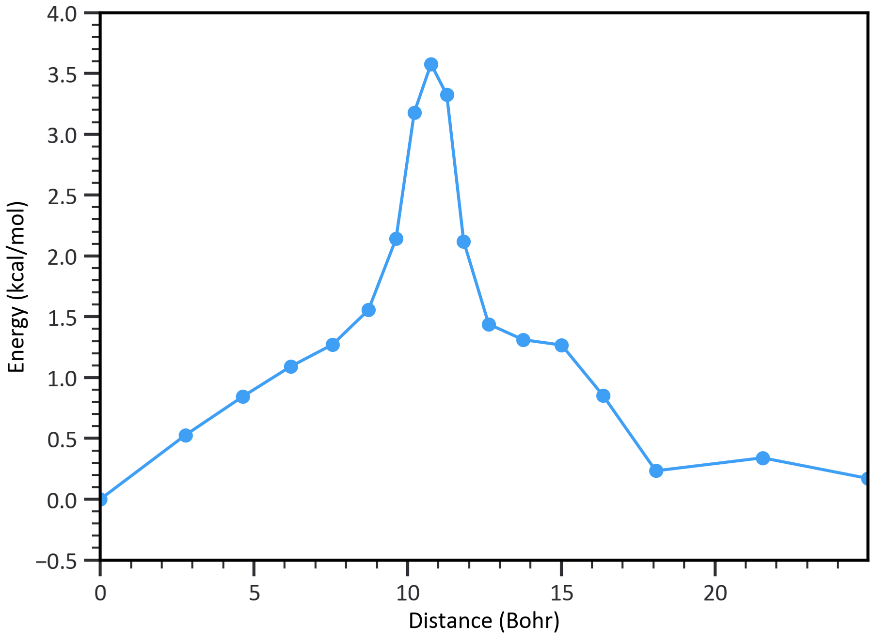
4.0
3.5
3.0
2.5
2.0
1.5
1.0
0.5
0.0
−0.5
Energy (kcal/mol)
0
5
10
15
20
Distance (Bohr)

**f**

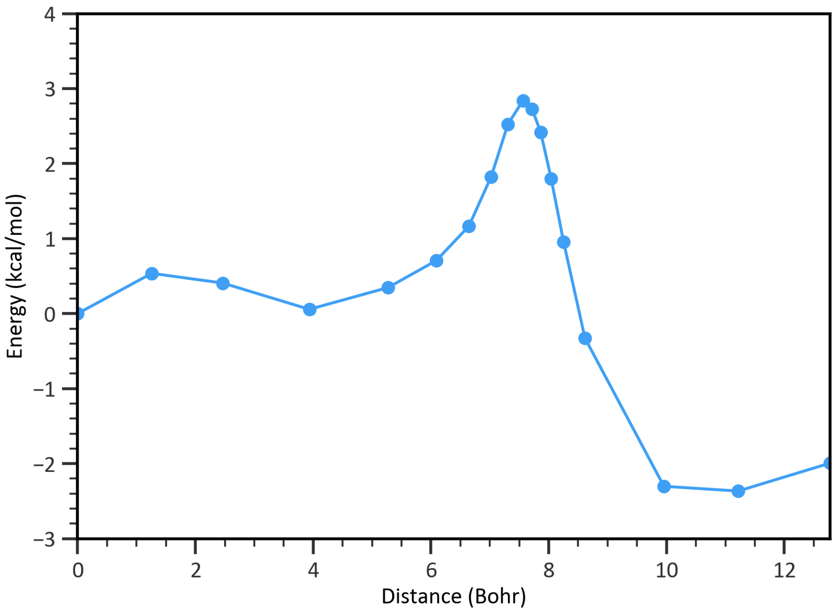
4
3
2
1
0
-1
−2
−3
Energy (kcal/mol)
0
2
4
6
8
10
12
Distance (Bohr)

**g**

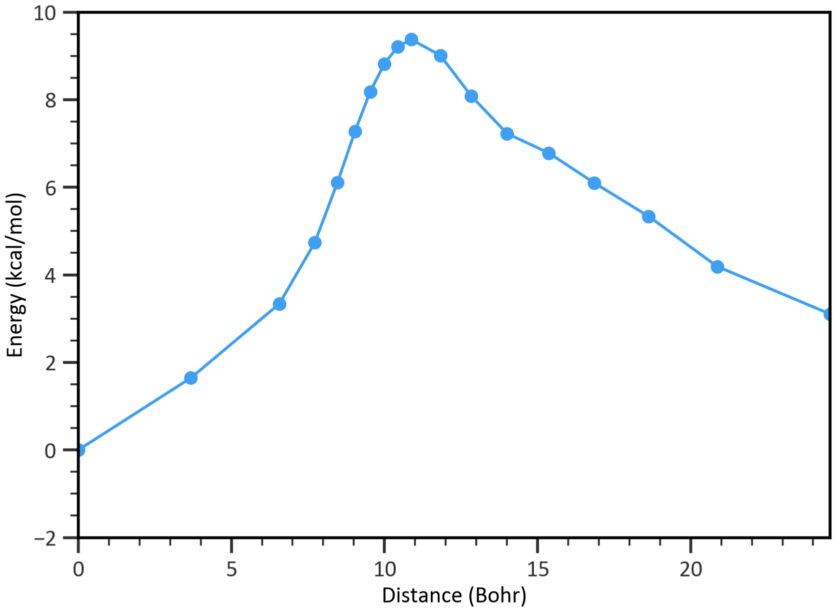


**h**

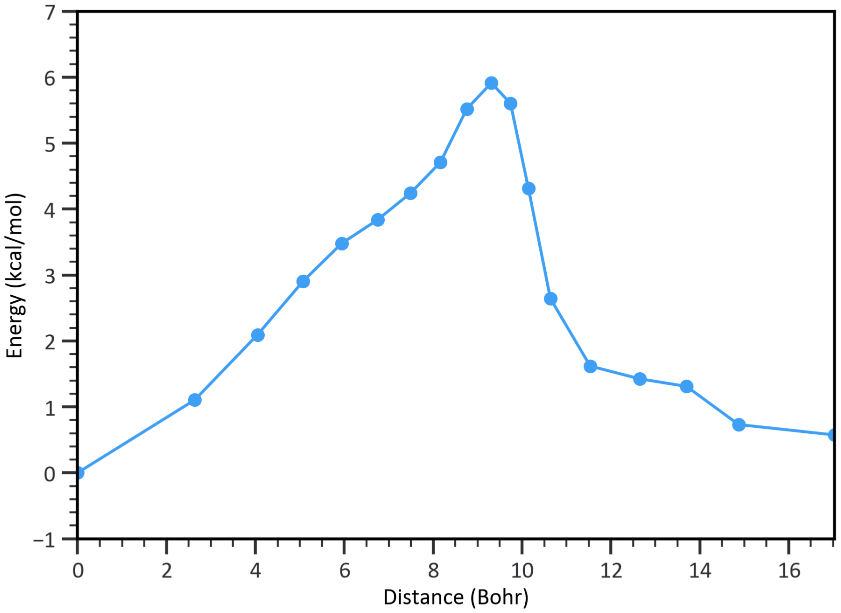


**Supplementary Fig. S4:** MEP plots of the chemical steps in the ZmTPS42/KSL1-catalyzed mechanism, as obtained from RxnNet. **a**, MEP plot for the **I1** → **I1'** step of the ZmTPS42/KSL1-catalyzed reaction mechanism generated using RxnNet. **b**, MEP plot for the **I1'** → **I1$^{NA}$** step. **c**, MEP plot for the **I1$^{NA}$** → **I2** step. **d**, MEP plot for the **I2** → **I2$^{NA}$** step. **e**, MEP plot for the **I2$^{NA}$** → **I3** step. **f**, MEP plot for the **I3** → **I4** step. **g**, MEP plot for the **I4** → **I4$^{NA}$** step. **h**, MEP plot for the **I4$^{NA}$** → **I5** step.
