## Supplementary Fig. S5: NMR structural elucidation of compound 13, epoxyrosanol. for "Discovery of the rosalexin pathway expands the modular network of maize diterpenoid chemical defenses"

**a**

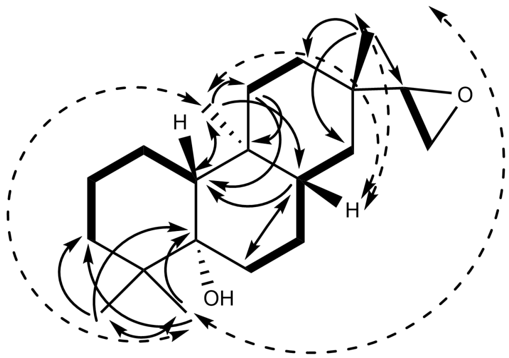


**b**

| Position | δC (ppm) | δH (ppm) |
|---|---|---|
| 1 a<br>b | 21.1785 | 1.626<br>1.358 |
| 2 a<br>b | 22.0695 | 1.589<br>1.477 |
| 3 a<br>b | 36.8689 | 1.665<br>1.121 |
| 4 | 38.8302 | |
| 5 | 76.6062 | |
| 6 a<br>b | 32.5735 | 1.722<br>1.572 |
| 7 a<br>b | 25.0819 | 1.524<br>1.121 |
| 8 | 41.1534 | 1.2869 |
| 9 | 36.7478 | |
| 10 | 48.5204 | 1.338 |
| 11 a<br>b | 34.5873 | 1.585<br>1.111 |
| 12 a<br>b | 29.3628 | 1.578<br>1.204 |
| 13 | 33.365 | |
| 14 a<br>b | 34.4213 | 1.422<br>1.003 |
| 15 | 61.1037 | 5.831 |
| 16 a<br>b | 43.4778 | 4.934<br>4.859 |
| 17 | 20.4828 | 1.0507 |
| 18 | 24.3954 | 0.884 |
| 19 | 23.977 | 1.0342 |
| 20 | 11.9278 | 0.919 |

c

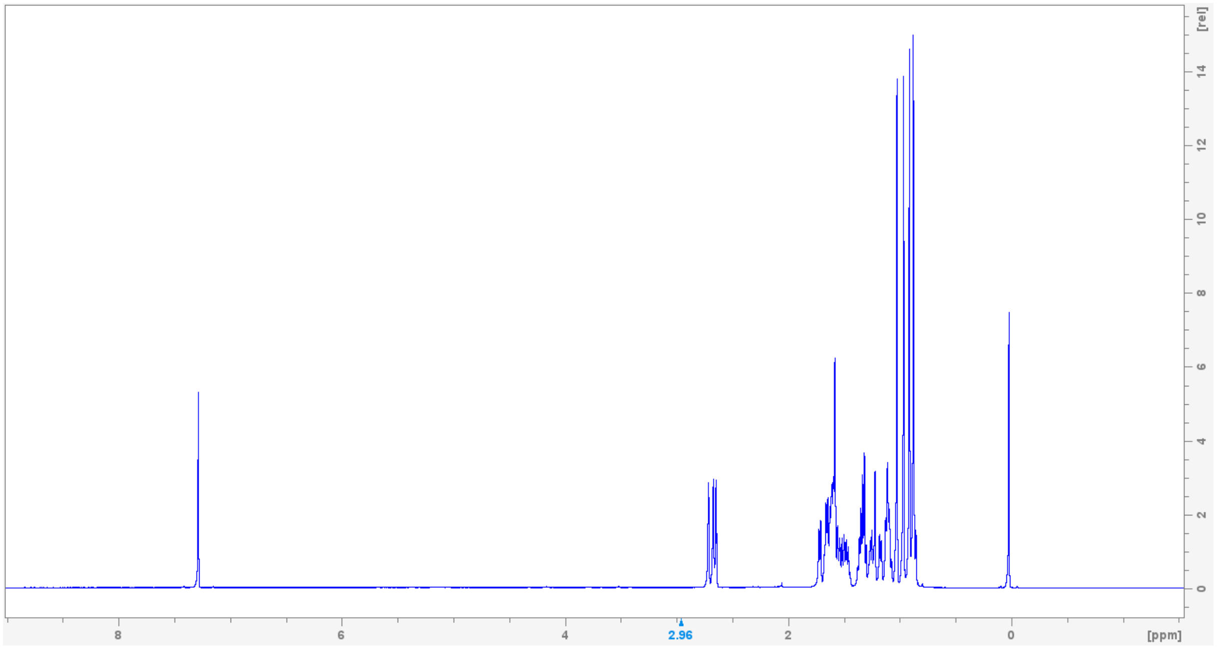
[rel]
14
12
10
8
6
4
2
0
8
6
4
2.96
2
0
[ppm]

d

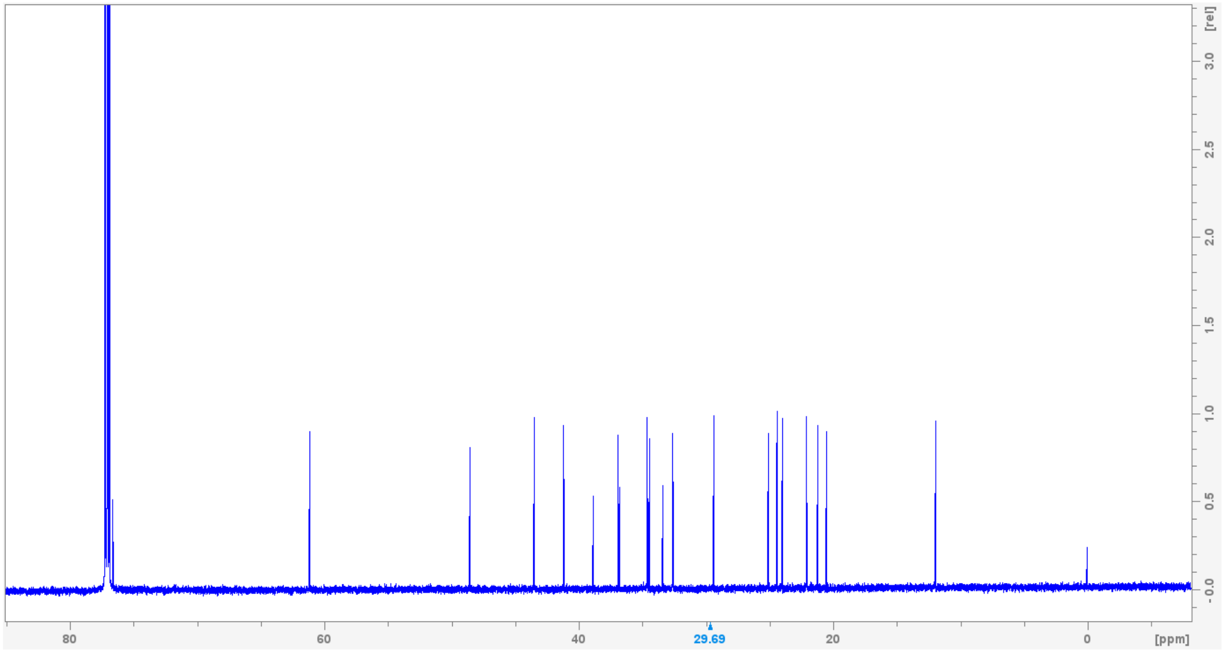
[rel]
3.0
2.5
2.0
1.5
1.0
0.5
-0.0
80
60
40
29.69
20
0
[ppm]

e

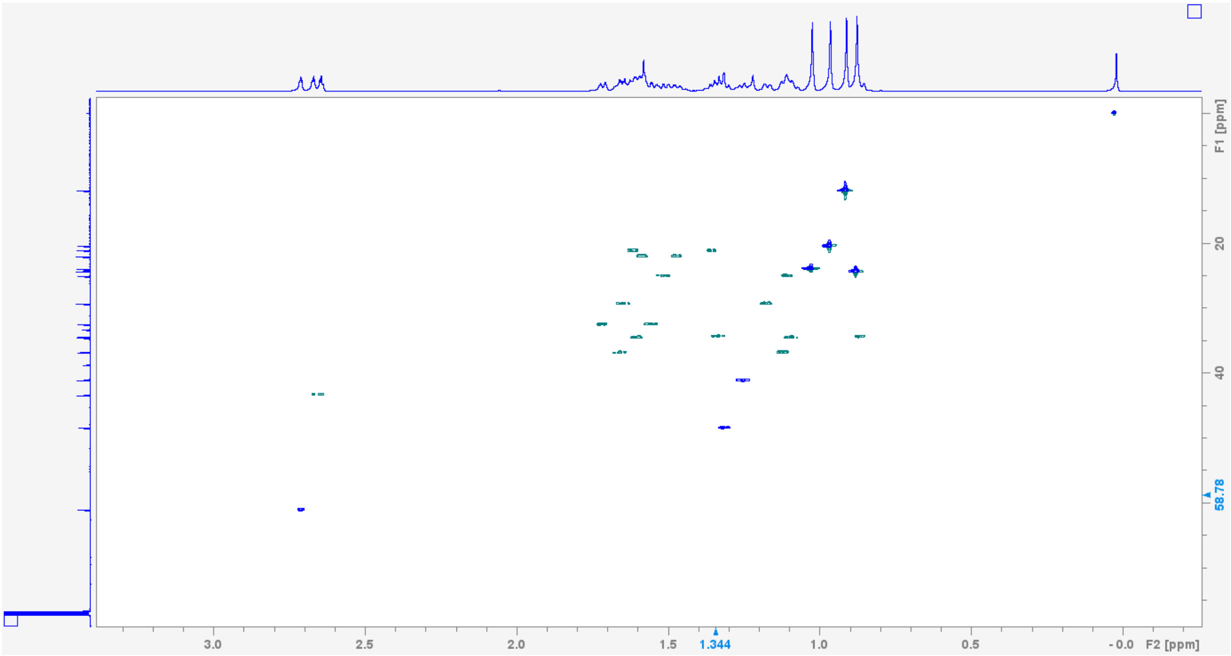
F1 [ppm]
20
40
58.78
3.0
2.5
2.0
1.5
1.344
1.0
0.5
- 0.0
F2 [ppm]

f

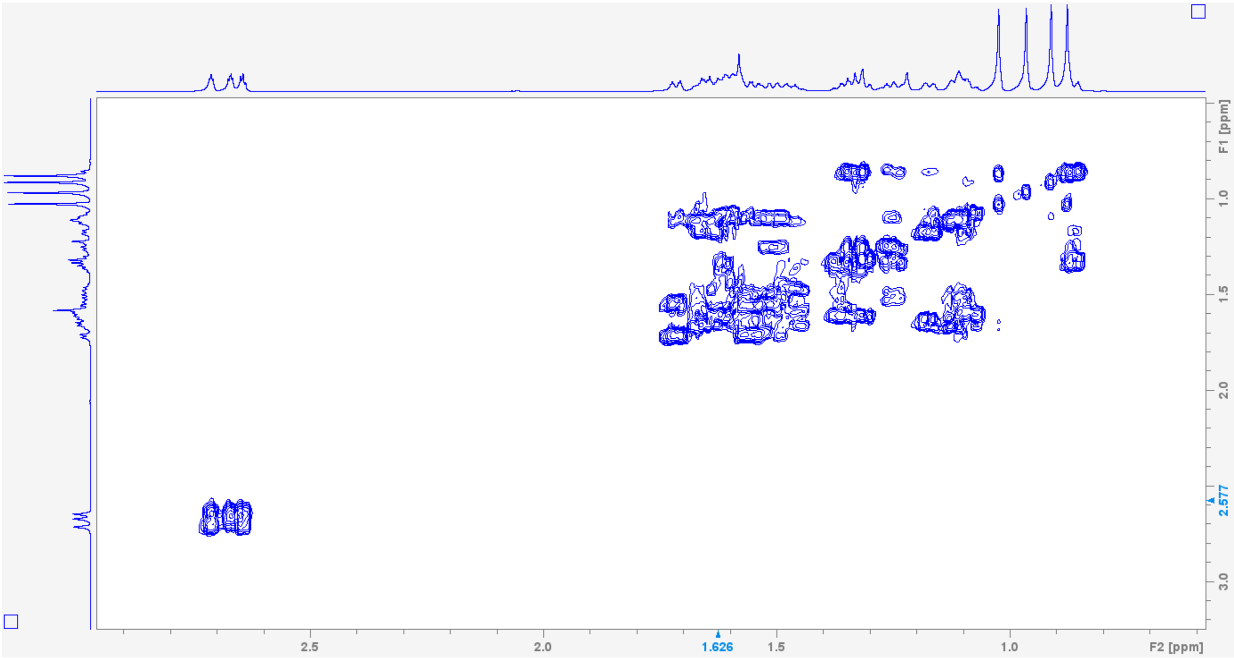
F1 [ppm]
1.0
1.5
2.0
2.577
3.0
2.5
2.0
1.626
1.5
1.0
F2 [ppm]

g

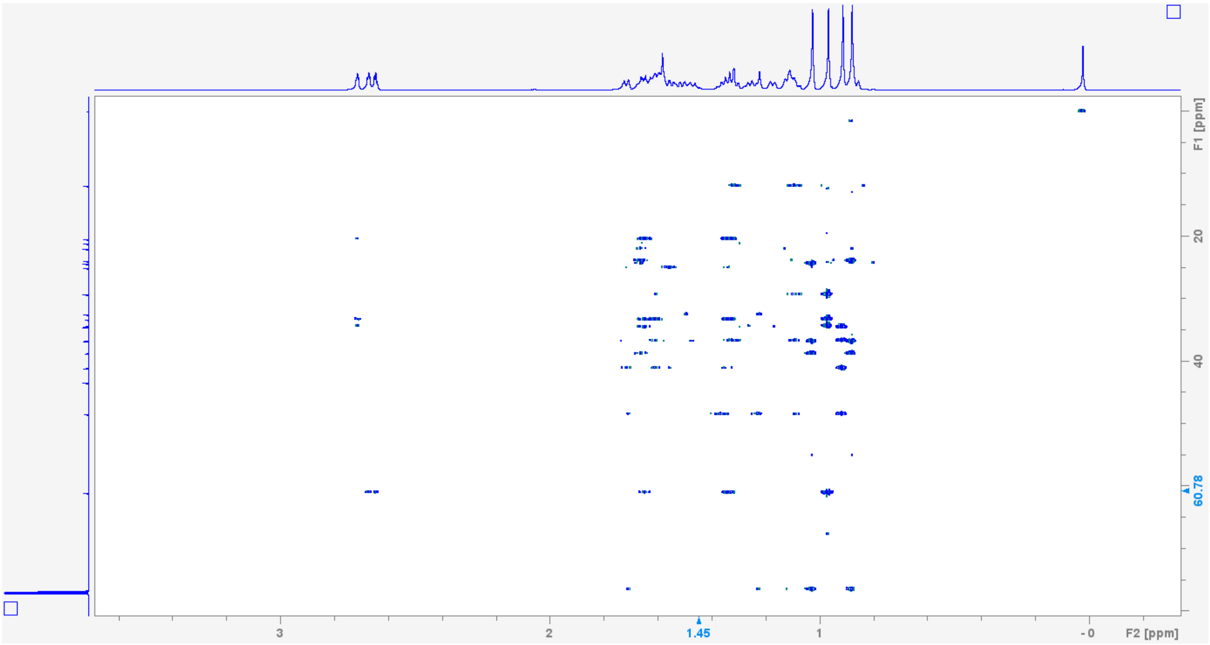


h

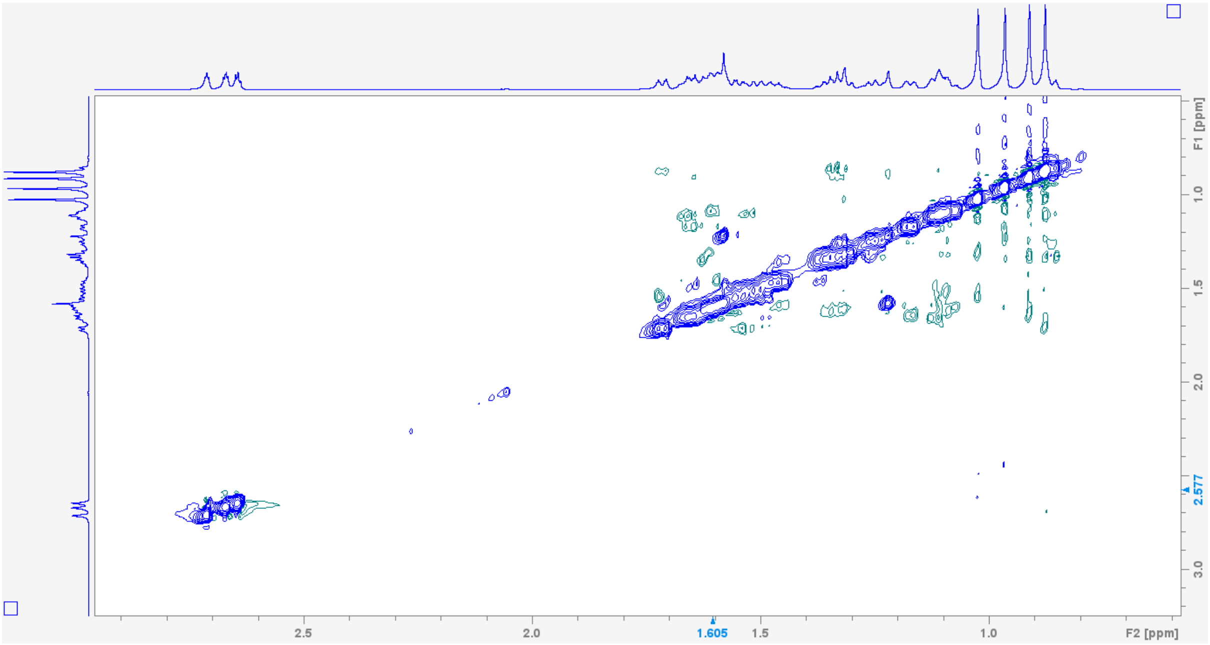


**Supplementary Figure S5.** NMR structural elucidation of compound **13**, epoxyrosanol. **a,** chemical structure depicted with key structural elements solved by experimental NMR including COSY (bold) and HMBC (arrows). **b**, Epoxyrosanol $^{13}$C and $^{1}$H assignments and chemical shift table. Epoxyrosanol NMR spectra: **c**, $^{1}$H NMR; **d**, $^{13}$C NMR; **e**, HSQC; **f**, COSY; **g**, HMBC; **h**, NOESY.
