## Supplementary Fig. S6: NMR structural elucidation of compound 14, epoxyrosaene. for "Discovery of the rosalexin pathway expands the modular network of maize diterpenoid chemical defenses"

**a**

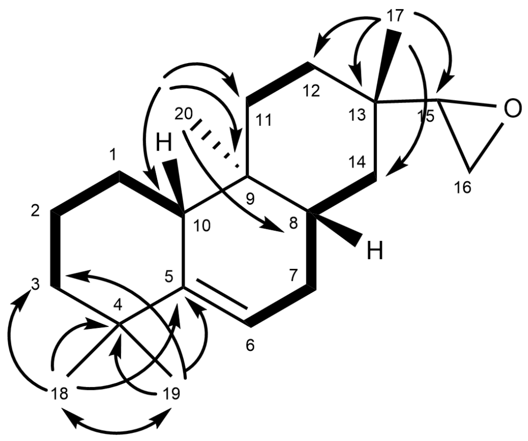


**b**

| Position | δC (ppm) | δH (ppm) |
|---|---|---|
| 1 a<br>b | 26.5654 | 1.802<br>2.093 |
| 2 a<br>b | 21.9358 | 1.595<br>1.539 |
| 3 a<br>b | 40.8027 | 1.412<br>1.237 |
| 4 | 35.8912 | |
| 5 | 145.8443 | |
| 6 | 116.2253 | 5.492 |
| 7 a<br>b | 30.2062 | 2.19<br>1.677 |
| 8 | 35.6569 | 1.455 |
| 9 | 34.8835 | |
| 10 | 47.1978 | 1.923 |
| 11 a<br>b | 33.7430 | 1.698<br>1.270 |
| 12 a<br>b | 29.8558 | 1.560<br>1.231 |
| 13 | 33.3388 | |
| 14 a<br>b | 34.2599 | 1.222<br>1.010 |
| 15 | 61.2331 | 2.72 |
| 16 a<br>b | 43.3800 | 2.678<br>2.644 |
| 17 | 19.5772 | 0.946 |
| 18 | 29.3412 | 1.029 |
| 19 | 29.7230 | 1.081 |
| 20 | 12.3750 | 0.674 |

c

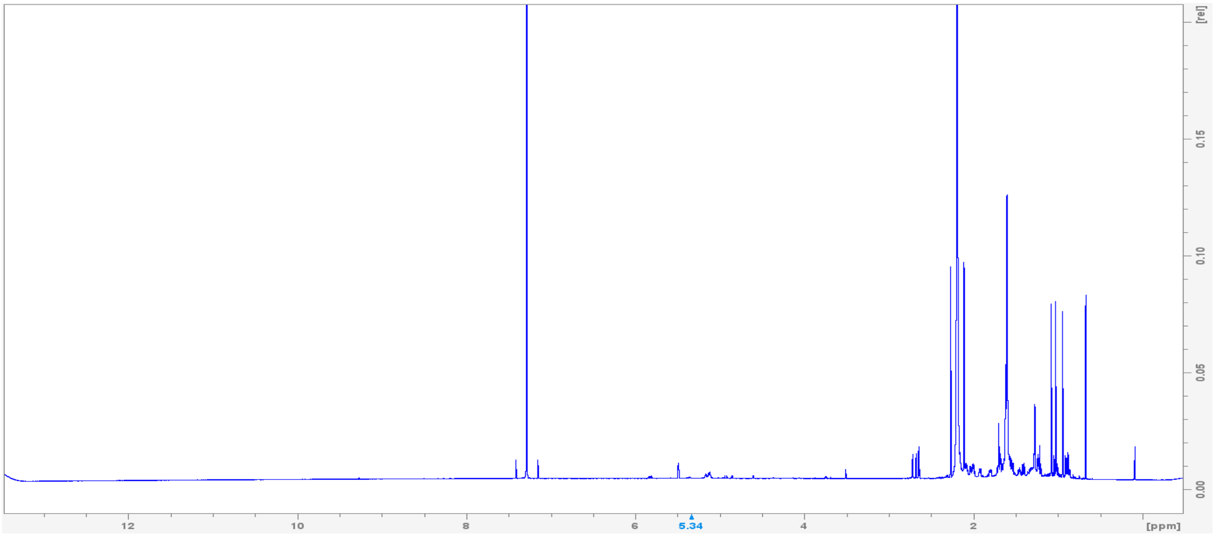
[rel]
0.15
0.10
0.05
0.00
12
10
8
6
5.34
4
2
[ppm]

d

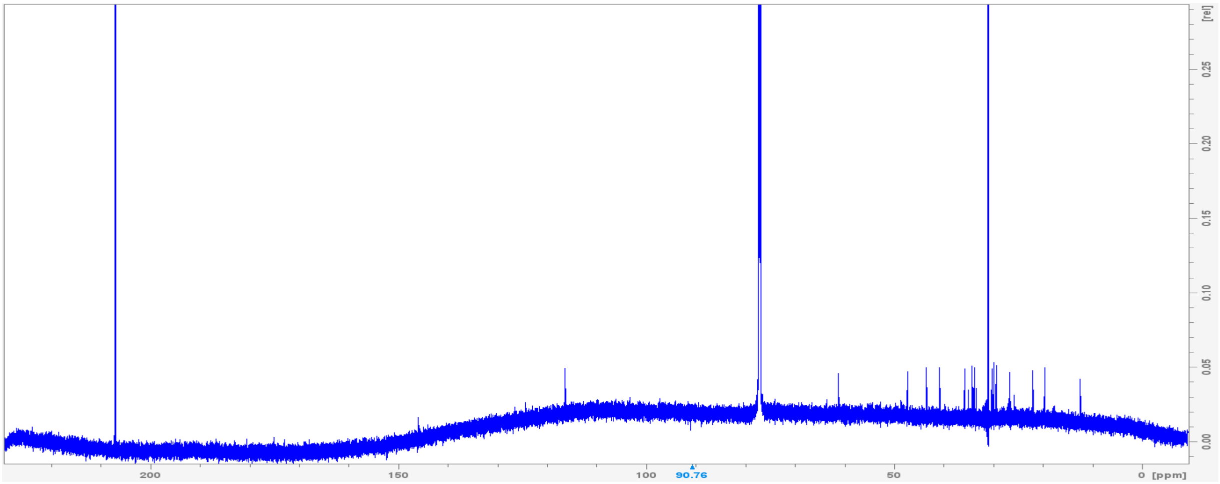
[rel]
0.25
0.20
0.15
0.10
0.05
0.00
200
150
100
90.76
50
0
[ppm]

e

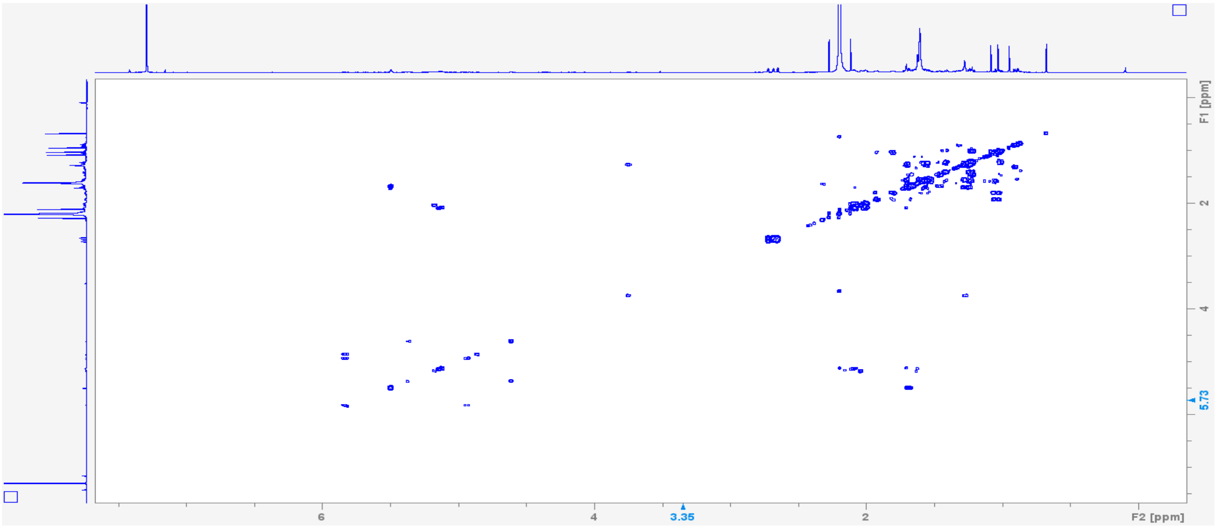
F1 [ppm]
2
4
5.73
6
4
3.35
2
F2 [ppm]

f

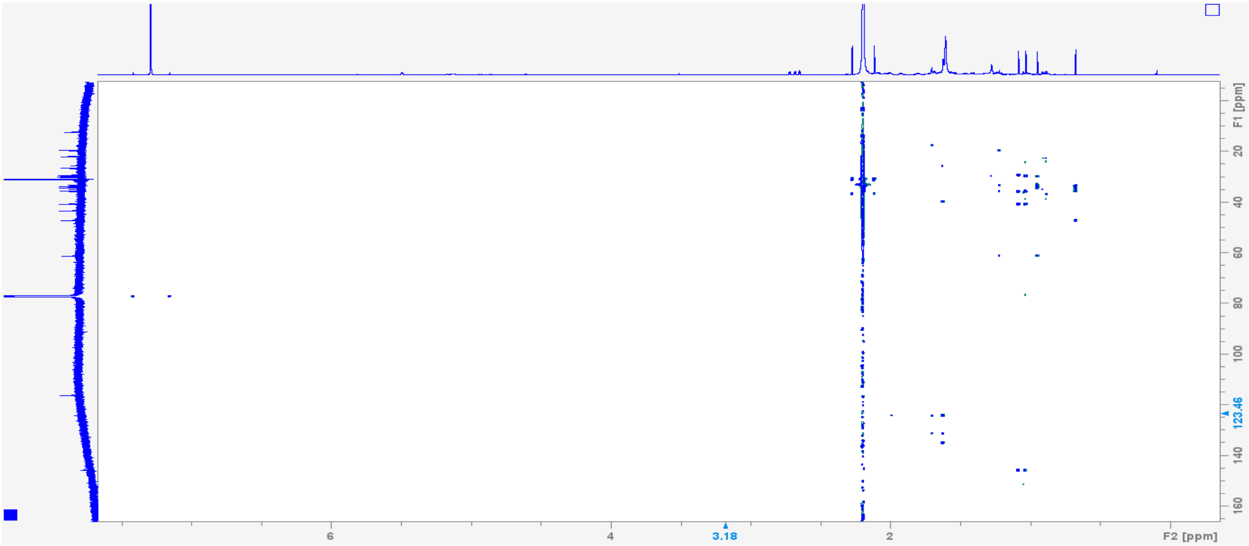
F1 [ppm]
20
40
60
80
100
123.46
140
160
6
4
3.18
2
F2 [ppm]

g

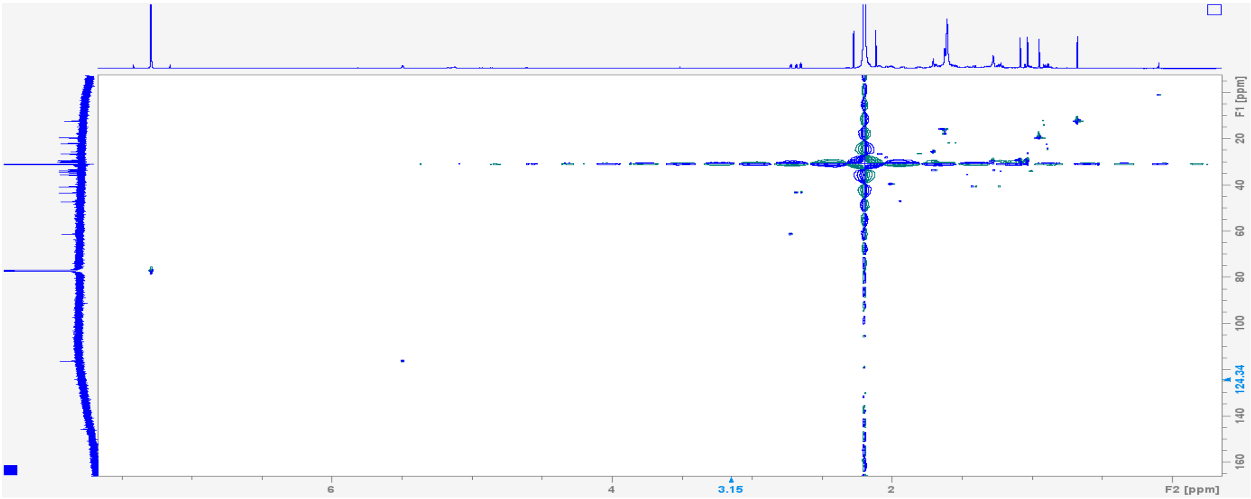
F1 [ppm]
20
40
60
80
100
124.34
140
160
6
4
3.15
2
F2 [ppm]

h

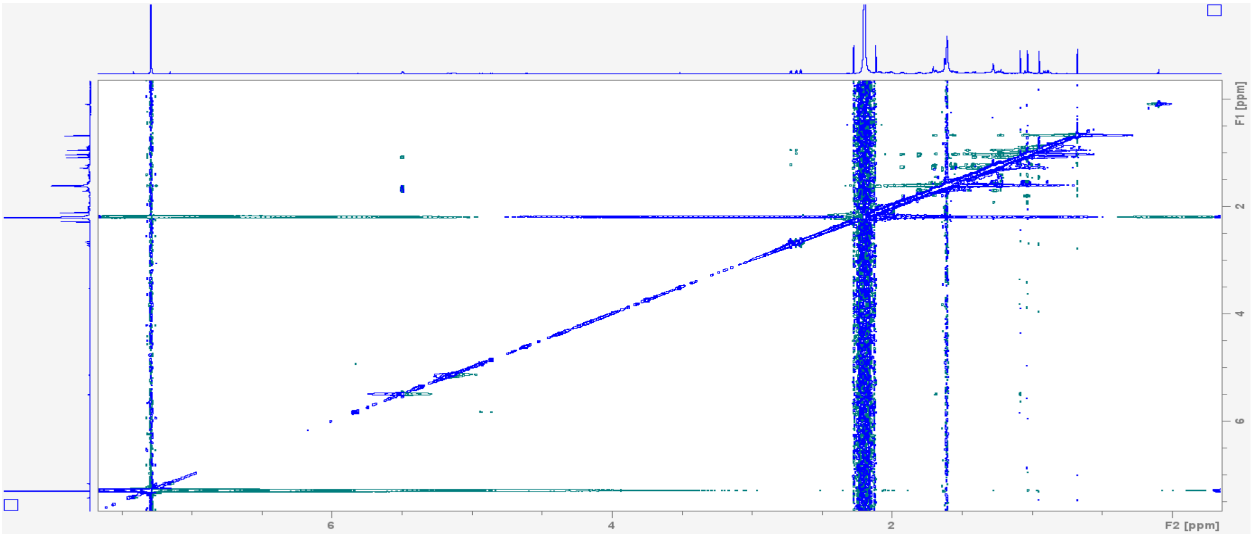
F1 [ppm]
2
4
6
6
4
2
F2 [ppm]

**Supplementary Figure S6.** NMR structural elucidation of compound **14**, epoxyrosaene. **a**, chemical structure depicted with key structural elements solved by experimental NMR including COSY (bold) and HMBC (arrows). **b**, Epoxyrosaene 13C and 1H assignments and chemical shift table. Epoxyrosaene NMR spectra: **c**, 1H NMR; **d**, 13C NMR; **e**, HSQC; **f**, COSY; **g**, HMBC; **h**, NOESY.
