## Supplementary Fig. S7: Functional analysis of maize P450 enzymes for 5-rosanol conversion. for "Discovery of the rosalexin pathway expands the modular network of maize diterpenoid chemical defenses"

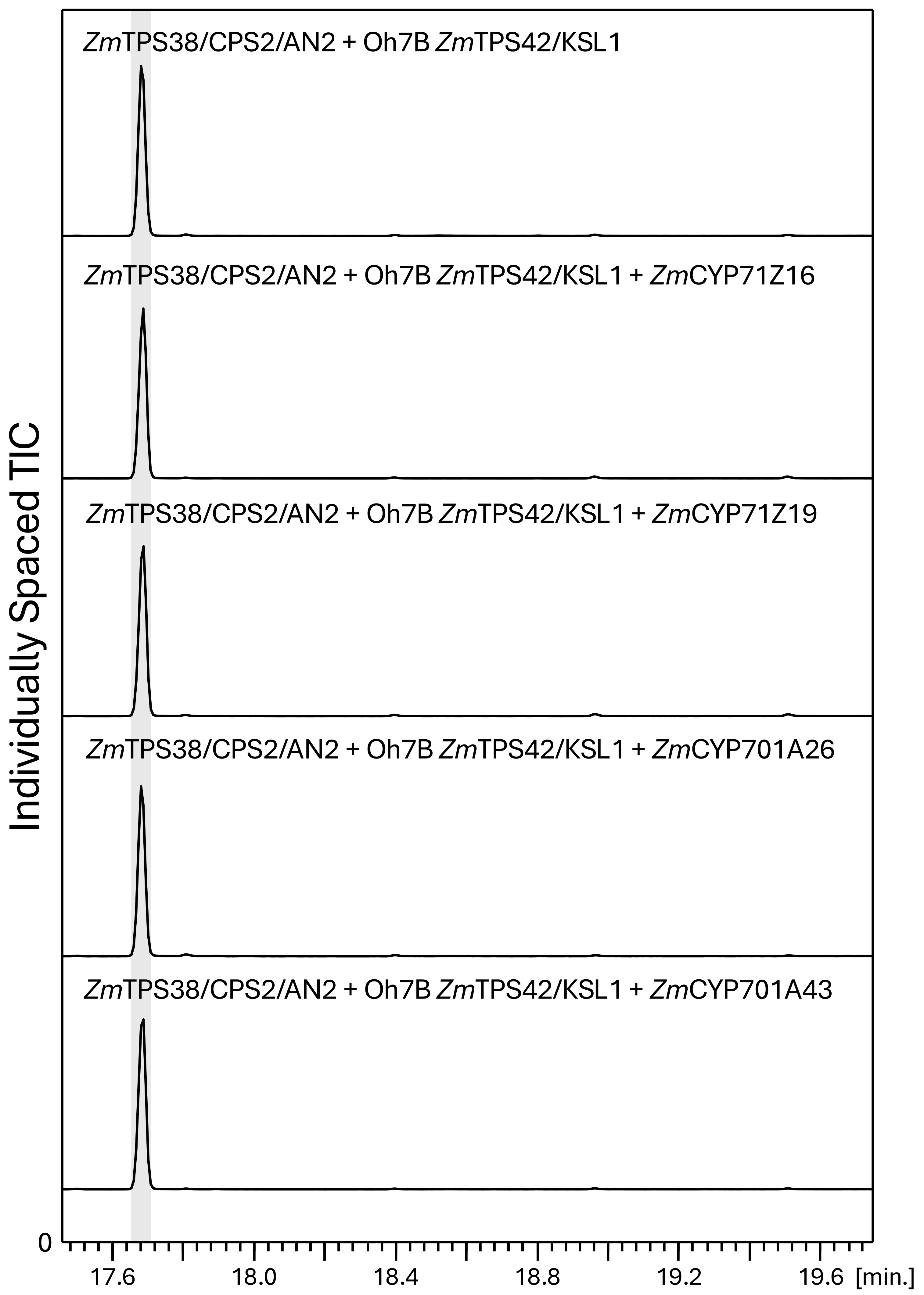


**Supplementary Fig. S7 |** Functional analysis of maize P450 enzymes for 5-rosanol conversion. Individually scaled total ion chromatograms (TIC) of reaction products resulting from *Nicotiana benthamiana* co-expression assays of ZmTPS38CPS2/AN2 and ZmTPS42/KSL1 only, or with the addition of known maize diterpenoid-metabolic P450 enzymes, ZmCYP71Z16, ZmCYP71Z19, ZmCYP701A26, and ZmCYP701A43. Highlighted peak depicts 5-rosanol.
