## Supplementary Fig. S8: NMR structural elucidation of compound 15, trihydroxyrosanol (THR). for "Discovery of the rosalexin pathway expands the modular network of maize diterpenoid chemical defenses"

**a**

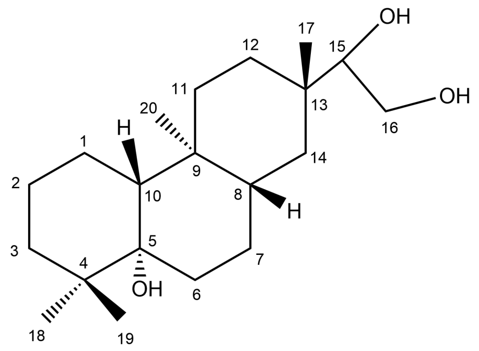


**b**

| Position | δC (ppm) | δH (ppm) |
|---|---|---|
| 1 a<br>b | 20.6 | 1.578<br>1.469 |
| 2 a<br>b | 21.94 | 1.567<br>1.506 |
| 3 a<br>b | 36.49 | 1.749<br>1.063 |
| 4 | 38.49 | |
| 5 | 76.45 | |
| 6 a<br>b | 31.94 | 1.778<br>1.571 |
| 7 a<br>b | 25.13 | 1.571<br>1.092 |
| 8 | 41.21 | 1.251 |
| 9 | 36.41 | |
| 10 | 48.69 | 1.31 |
| 11 a<br>b | 34.7 | 1.596<br>1.119 |
| 12 a<br>b | 28.99 | 1.683<br>1.083 |
| 13 | 36.39 | |
| 14 a<br>b | 35.79 | 1.412<br>1.093 |
| 15 | 81.71 | 3.2 |
| 16 a<br>b | 62.15 | 3.45<br>3.72 |
| 17 | 18.39 | 0.935 |
| 18 | 23.75 | 0.863 |
| 19 | 23.2 | 1.026 |
| 20 | 11.14 | 0.945 |

c

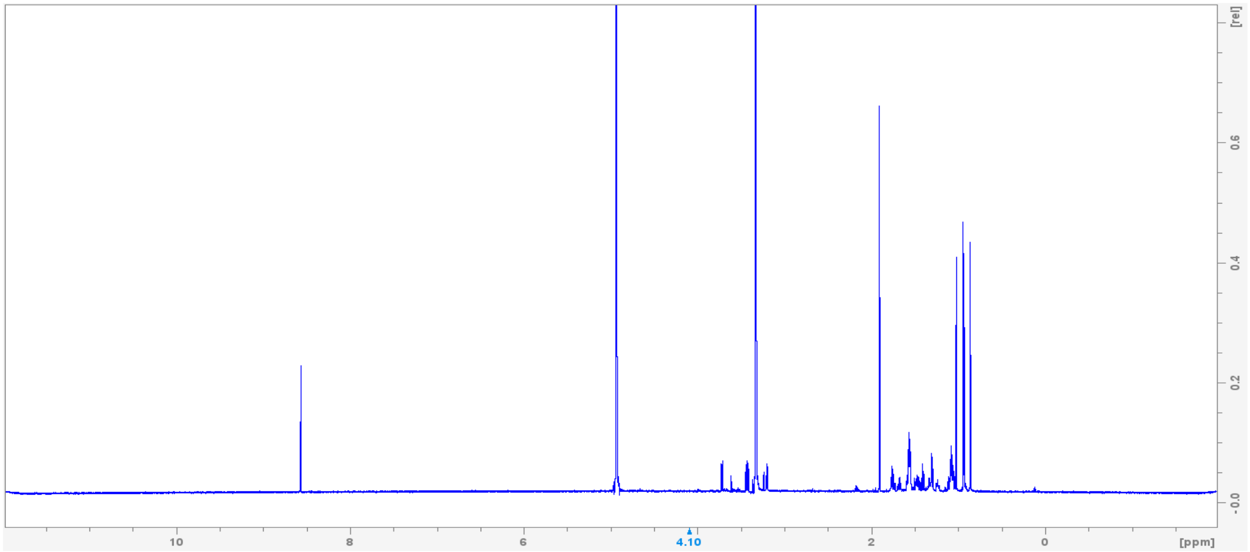
[rel]
0.6
0.4
0.2
-0.0
10
8
6
4.10
2
0
[ppm]

d

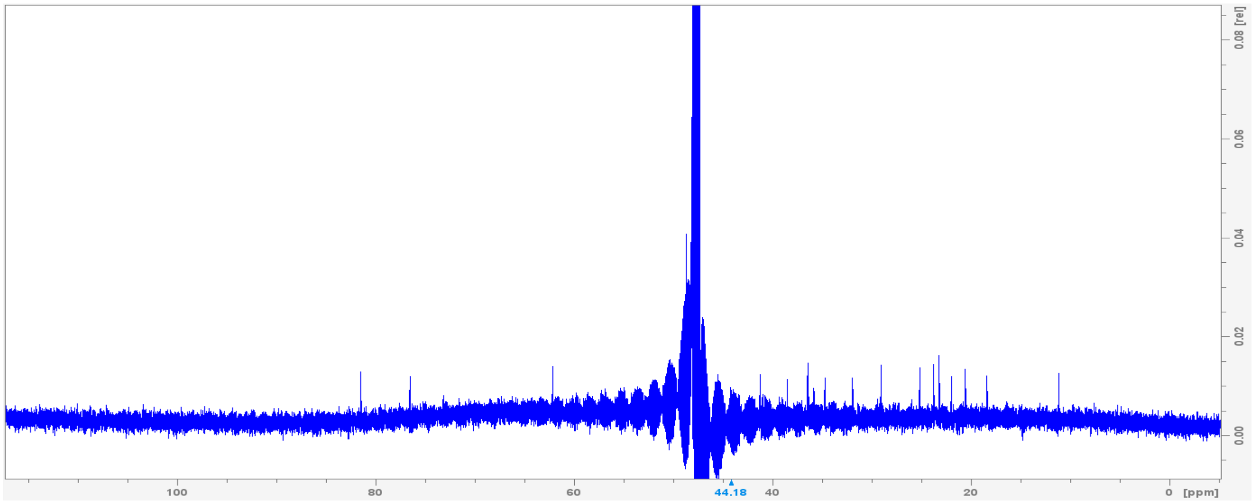
0.08 [rel]
0.06
0.04
0.02
0.00
100
80
60
44.18
40
20
0
[ppm]

e

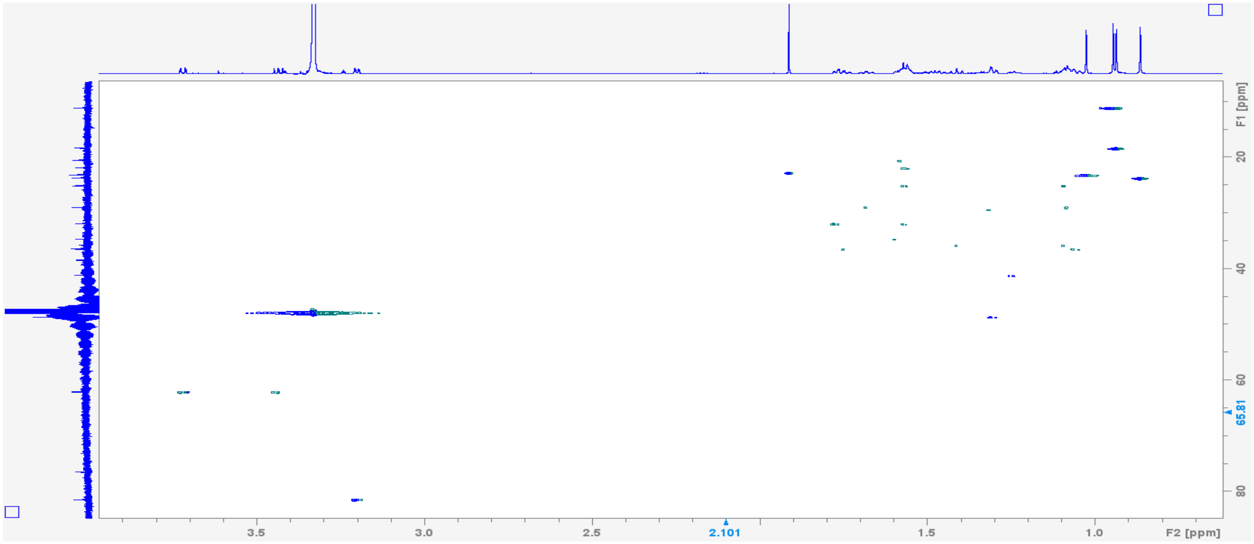
F1 [ppm]
20
40
60
65.81
80
3.5
3.0
2.5
2.101
1.5
1.0
F2 [ppm]

**Supplementary Figure S8.** NMR structural elucidation of compound **15**, trihydroxyrosanol (THR). **a**, chemical structure. **b**, Trihydroxyrosanol 13C and 1H assignments and chemical shift table. Trihydroxyrosanol NMR spectra: **c**, 1H NMR; **d**, 13C NMR; **e**, HSQC.
